# Decoding microbiome and protein family linkage to improve protein structure prediction

**DOI:** 10.1101/2021.04.15.440088

**Authors:** Pengshuo Yang, Wei Zheng, Kang Ning, Yang Zhang

## Abstract

Information extracted from microbiome sequences through deep-learning techniques can significantly improve protein structure and function modeling. However, the model training and metagenome search were largely blind with low efficiency. Built on 4.25 billion microbiome sequences from four major biomes (Gut, Lake, Soil and Fermentor), we proposed a MetaSource model to decode the inherent link of microbial niches with protein homologous families. Large-scale protein family folding experiments showed that a targeted approach using predicted biomes significantly outperform combined metagenome datasets in both speed of MSA collection and accuracy of deep-learning structure assembly. These results revealed the important link of biomes with protein families and provided a useful bluebook to guide future microbiome sequence database and modeling development for protein structure and function prediction.

---

Given the rapid explosion of protein sequences, computer-based approaches play an increasingly important role in protein structure determination and structure-based function annotations (*1, 2*). Two types of strategies have been widely considered for protein 3D structure prediction (*2*): the first is template-based modeling (TBM), which constructs structural models using solved structures as templates, where its success requests for the availability of homologous templates in the Protein Data Bank (PDB); the second is template-free modeling (FM) approach (or *ab initio* modeling), which dedicates to model the “Hard” proteins that do not have close homologous structures in the PDB. Due to the lack of reliable physics-based force fields, the most efficient FM methods, including Rosetta (*3*), QUARK (*4*), and I-TASSER (*5*), rely on *a prior* spatial restraints derived, usually through deep neural-network learning (*6, 7*), from the co-evolution information based on multiple sequence alignments (MSA) of homologous proteins (*8*). Hence, to model 3D structure of the “Hard” proteins, a sufficient number of homologous sequences is critical to ensure the accuracy of deep machine-learning models and the quality of subsequent 3D structure constructions (*9*).

Considerable effort was recently paid to the utilization of metagenome sequence data to enhance the MSA and FM model constructions. For example, Ovchinnikov *et al*. used the Integrated Microbial Genomes (IMG) database to generate contact-map predictions and create high-confidence models for 614 Pfam protein families that lack homologous structures in the PDB (*10*). Using UniRef20 (*11*), Michel *et al*. combined contact-map prediction with the CNS folding method (*12*) to model protein structure for 558 Pfam families of unknown structure with an estimated 90% specificity. Most recently, Wang *et al*. examined the usefulness of the *Tara* Oceans microbial genomes and found that the microbiome genomes can provide additional help on high-quality MSA construction and protein structure and function modeling (*13*). This result demonstrated a significant role of the microbiome sequences, which represent one of the largest reservoirs of microbial species on this planet, in FM structural folding and structure-based function annotations.

Despite the success of metagenome-assisted 3D structure modeling, there are still thousands of Pfam families whose structure cannot be appropriately modeled with a satisfactory confidence. One critical reason is that despite the rapid accumulation of sequences, the current sequence databases are far from complete and very few homologous sequences are available for many of the FM targets. On the other hand, the metagenome sequence databases have become extremely large (e.g., the JGI database contains more than 60 billion microbial genes) (*14, 15*), which makes a thorough and balanced database search increasingly slow and difficult. In a recent study, Zhang *et al*. showed that using current data mining tools, the quality of MSAs from metagenome library is not always proportional to the effective number of homologous sequences (*Neff*, see **Eq. S3** in **Supplemental Materials**), partly due to the complexity of the sequence family relations and the bias of sequence database searches (*8*). The recent CASP experiments also witnessed various examples where the folding simulations for FM targets are negatively impacted by the contact/distance predictions due to the biased MSAs from the large metagenome datasets despite the high *Neff* value (*16, 17*). Therefore, a deeper understanding of the inherent links between the metagenome and protein families, if exist, should be of critical importance to help improve the efficiency of sequence database searching and the subsequent 3D structure modeling.

In this work, we aim to decipher the relationship between microbial niches (biome), sequence and Pfam homology families through large-scale sequence-based structure folding analyses. We first collected 4.25 billion microbiome sequences from the EBI metagenomic database (MGnify database) (*18*) that cover four major biomes (Gut, Lake, Soil and Fermentor), to examine their capacities for assisting 3D protein structure prediction on unsolved Pfam families. The “marginal effect” analyses showed profoundly different effects of specific biomes on supplementing homologous sequences for different Pfam families. A machine learning model named MetaSource is then developed to predict the source biome of target proteins, which can significantly improve the speed and memory request of MSA search and subsequent 3D structure modeling accuracy. These results have validated the important biome-sequence-Pfam associations, leading a novel way towards better efficiency and effectiveness of the microbiome-based targeted approach to protein structure and function predictions.

## Results and Discussions

### Biome-specific microbiome samples contain billions of different functional genes from thousands of genera

1,705 microbiome samples were collected from four typical microbial niches (biomes) (Gut, Lake, Soil and Fermentor, **Figure 1A**). Processed by the EBI pipeline version 4.1, a total of 4.25 billion protein sequences (functional genes) were predicted from these biomes, where a biome-specific taxonomic profile can be observed in **Figure 1B**.

**Figure 1.**
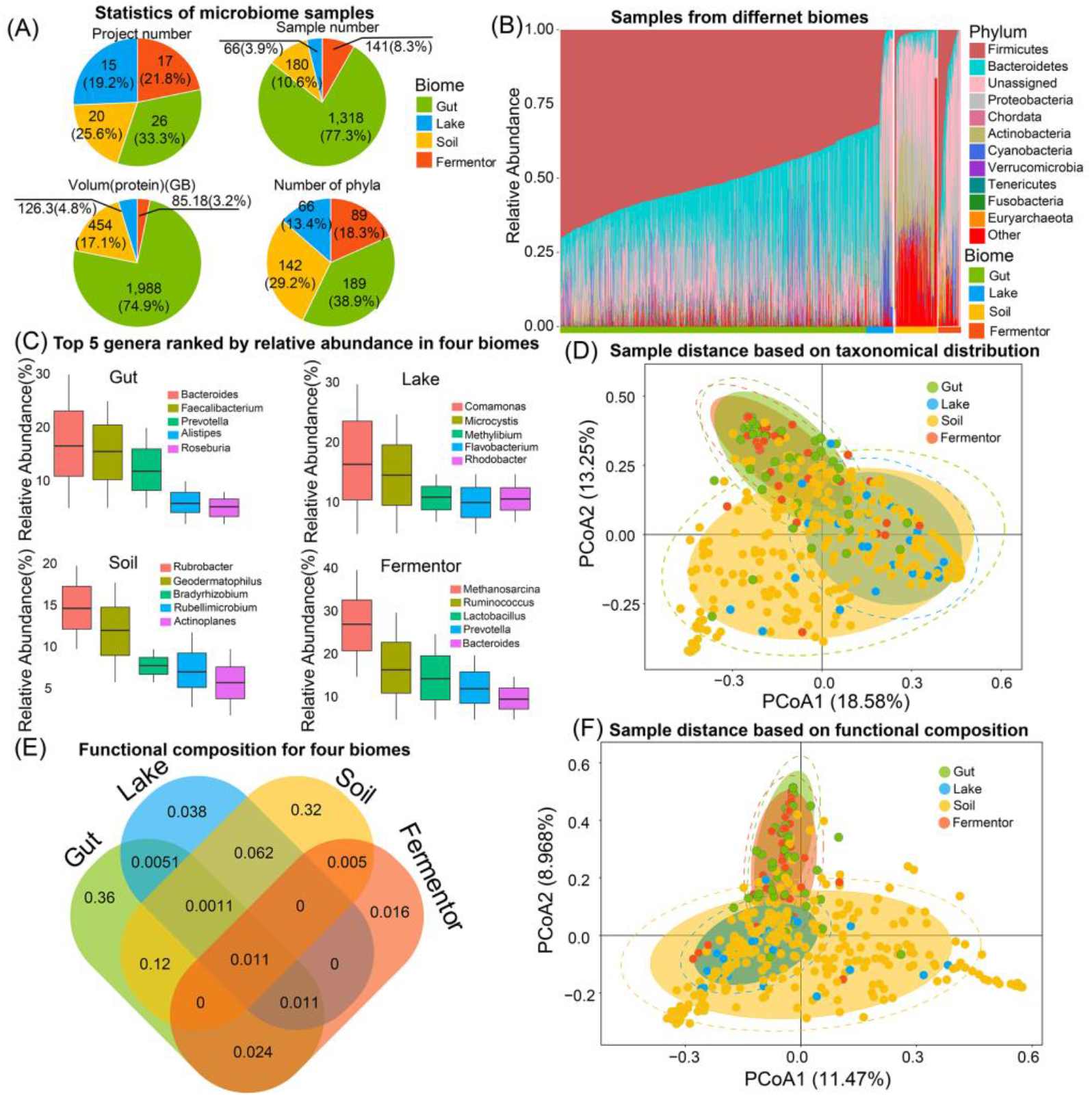
Taxonomic and functional profiling for different microbiome samples. (A) The basic statistics of microbiome samples collected from the four biomes. (B) Species distribution on phylum level for samples in four biomes. The species distribution is categorized by their biomes and labeled with different colors. For all the samples, the top 10 phyla ranked by the average counts among all samples are illustrated. “Unassigned” means the species cannot be identified by a known phylum. “Other” represent the combination of rest phyla. (C) Top five genera ranked by relative abundances for four biomes. (D) PCoA result based on taxonomic profile on genus level for samples from the four biomes. Samples from the same biome are labeled with the same color. The confidence intervals of samples in the same biome are marked in circles. (E) The shared and specific functional distribution for four biomes. The number labeled in the figure means the number (in billion) of specific or sheared sequences annotated by GO database. (F) PCoA result based on functional distribution for samples from the four biomes based on GO annotation. Samples from the same biome are labeled with the same color. The confidence intervals of samples in the same biome are marked in circles.

Among the 1,705 microbiome samples, 169 phyla were identified, covering the common members in the kingdom of Bacteria and Archaea. Further classification on the genus level, 8,721 genera were identified, and different top five genera ranked by relative abundance in four biomes also illustrate a biome-specific taxonomic profile (**Figure 1C**). These results indicate that the biomes host different microbiome cohorts and further investigation revealed the correlation between microbial communities’ taxonomic profile and their living environment: In the Gut biome, for example, Firmicutes (average relative abundance: 0.41±0.28) and Bacteroidetes (average relative abundance: 0.26±0.14) were the dominant phyla. Members in phylum Firmicutes were involved in energy resorption associated with reduced low-grade inflammation in obesity (*19*). Bacteroidetes play an important role in the development of immune dysregulation and systemic disease (*20, 21*). In Lake and Soil biome, phylum Proteobacteria is the dominant phylum (average relative abundance: 0.23 ±0.18 and 0.35 ±0.16, respectively), which takes part in nitrogen fixation and oxidation of iron, sulfur, and methane (*22*). In the Fermentor biome, phylum Firmicutes is the dominating phylum (average relative abundance: 0.46 ± 0.36), in which most members play the role of anaerobic fermentation (*23*), the main function of most Fermentor.

To illustrate the divergence among the biomes, statistical tests were performed based on the species distribution: The Wilcox-test (nonparametric statistical test, single-tail test) for each pair of four biomes indicate a statistical difference among four biomes (**Table S1 in Supplementary Information, SI**). Furthermore, the Principal Coordinates Analysis (PCoA) indicates a biome-specific taxonomic profile for 1,705 microbiome samples (**Figure 1D**): samples collected from the same biome could cluster into one group (reflected by a concentrated confidence circle). Moreover, samples from the Lake biome were closer to those of the Soil biome, while those of the Gut biome and Fermentor biome were closer. This phenomenon could be attributed to the similar environments between Gut and Fermentor (oxygen-limited environment), as well as between Lake and Soil environment (open-air environment).

Among the 4.25 billion protein sequences obtained from these four biomes, we observed the biome-specific functional profiles. In total, 1.25 billion proteins could be annotated by GO database. Similar to taxonomic profile, these four biomes host different functional annotations (**Figure 1E**): 0.36 billion (68.4%) annotations were only detected in Gut biome, 0.038 (29.9%) billion annotations only in Lake biome, 0.32 (62.7%) billion annotations only in Soil biome and 0.016 billion (24.2%) of gene annotations only in Fermentor biome. The PCoA result based on functional profiles present clear differences among these four biomes (**Figure 1F**). Again, samples from Gut and Fermentor biomes were closer, while samples from Lake and Soil were closer, similar with the PCoA result based on taxonomic profiles.

### Metagenome-sourced proteins assisted successful structure modeling for thousands of protein families without homologous templates

Recent studies have shown that metagenome sequences can help improve the performance of protein structure prediction (*10, 13*), especially for the Pfam families without solved structures. Here, we selected 2,214 Pfam families with the *Neff* of MSA >16 from 8,700 Pfam families that have no member with solved structures. These Pfam families were all categorized as Hard targets by LOMETS because no homologous templates could be detected from the PDB by threading (*24*). We extended the contact-assisted I-TASSER method, C-I-TASSER (*17*), to predict structure models for the 2,214 unsolved Pfam families. Based on the benchmark results showing that models with a C-score >=-2.5 usually have a correct fold (see **Eq. S2** and **Figure S1**), 47% (1,044/2,214) of the Pfam families were found to be foldable by C-I-TASSER (**Figure 2A**). In **Figure 2B**, we presented the C-score histogram distribution of the C-I-TASSER models on the 2,214 unknown Pfam families. Considering false discovery rate (FDR) obtained from the benchmark tests (see **Materials and Methods**), there should be around 971 (=1,044*(1-6.96%)) Pfam families with high-confidence models.

**Figure 2.**
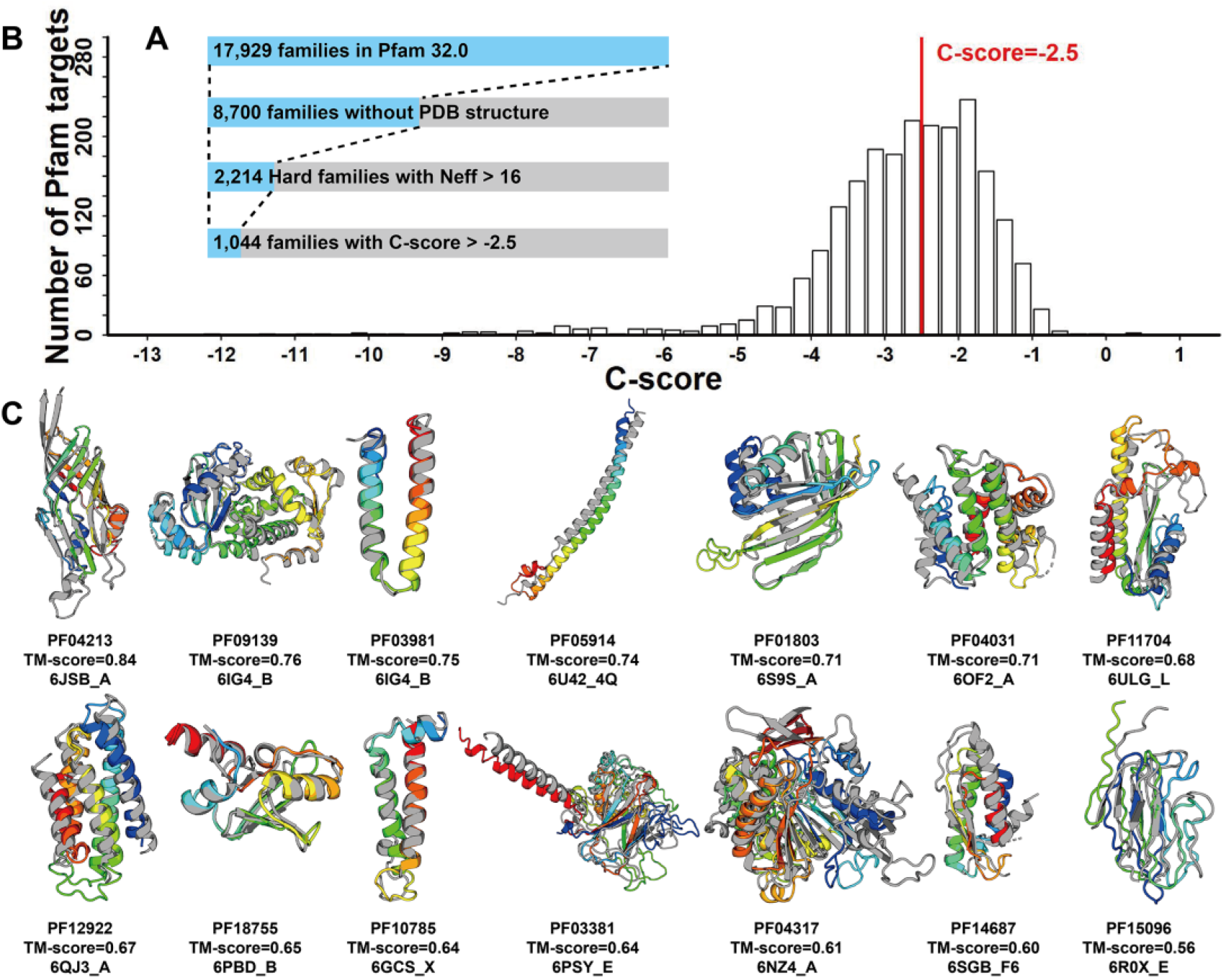
Structural modeling results for unknown Pfam Hard families. (A) Number of Pfam families at each stage of the analysis, where each set is a subset of the previous set. (B) The C-score distribution of the Pfam Hard families with *Neff* >16. (C) Structural models on 14 newly solved Pfam families with TM-score >0.5. In each case, the C-I-TASSER model is shown in rainbow color and the solved experimental structure of a member from the same Pfam family is shown in gray.

The C-I-TASSER modeling was performed on the Pfam database version 32.0 released in September 2018. The Pfam version 33.0 (released in March 2020) reported 28 new families with solved structures for at least one member among the 2,214 modeled Pfam families, which provides an opportunity to assess the performance of the prediction. Since only one member from each Pfam family was modeled by C-I-TASSER, the modeled member may be different from the member with solved structure. For these cases, we superposed the solved structure to the C-I-TASSER model using TM-align (*25*) and calculated the TM-score between the C-I-TASSER model and the experimental structure. The comparison between the C-I-TASSER models and the solved experimental structures is listed in **Table S2**. Although all the families are non-homologous to the PDB structures, 50% of the C-I-TASSER models have been correctly folded with TM-scores >0.5. This result is roughly consistent with the estimation that 47% of the 2,214 Pfam families are foldable by C-I-TASSER. **Figure 2D** presents the 14 Pfam families with successfully folded C-I-TASSER models, where C-I-TASSER models for all 2,214 unknown Pfam families are downloadable at https://github.com/HUST-NingKang-Lab/MetaSource.

### Enrichment of homologous sequences from different biomes

For the 1,044 Pfam families foldable by C-I-TASSER, an enrichment of homologous sequences from a specific biome can be observed, i.e., 964 Pfam families (964/1,044, 92.3%) could be identified with a single biome whose *Neff* value is larger than the other three biomes, including 105 families for Gut, 116 families for Lake, 617 families for Soil, and 126 for Fermentor (**Figure 3A**). For the remaining 80 Pfam families, two or more biomes contributed equally, which may be caused by the limited number of metagenome sequences (average 8.3±3.1 metagenome sequences) aligned. We observed that sequences from the Soil biome could assist in folding more Pfam families than other biomes, i.e., 39.6% sequences in the Soil biome could be aligned to Pfam families, while only 33.1% for the Gut biome, 30.8% for the Lake biome, and 24.3% for the Fermentor biome. These results are understandable as the metagenome in the Soil biome has been shown to have the highest species richness and most functional genes among these four biomes (*26*). However, it is worth mentioning that though microbiome sequences from Soil biome could supplement more Pfam than sequences from other biomes, this is not a winner-take-all situation: other biomes still work better than Soil biome for specific Pfam families.

**Figure 3.**
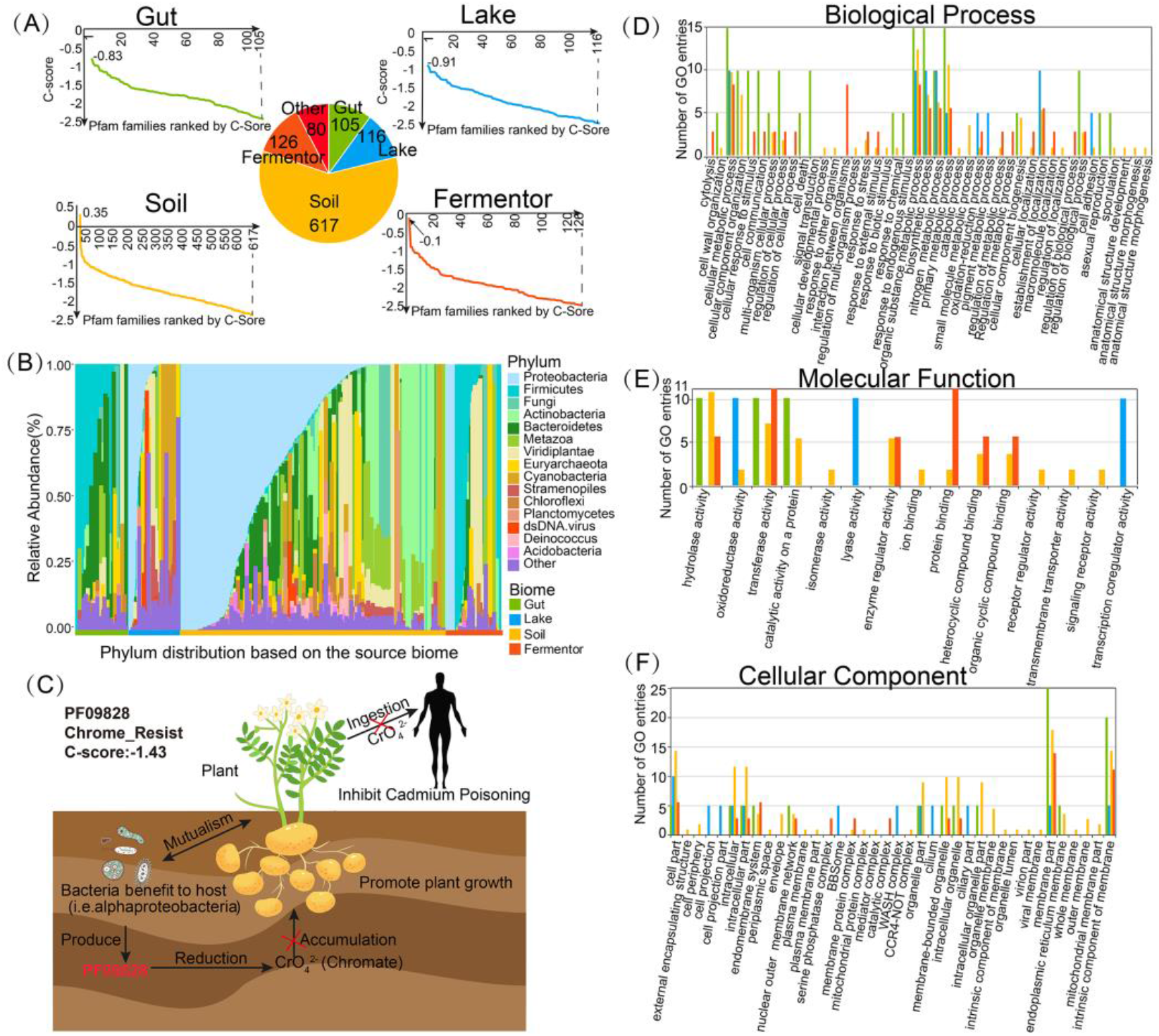
The taxonomic and functional properties of the Pfam families foldable by C-I-TASSER. (A) C-score distribution for Pfam families after replenishing by metagenome sequences. The vertical axis represents the C-score. For each panel, horizontal axis represents the Pfam families, (*25*). (B) The relative abundance of species distribution for Pfam families which were foldable by C-I-TASSER. The species distribution is divided into four biomes and labeled with different colors. Calculated by the average count among all samples, the top 10 phyla are illustrated and ranked. “Other” represent the combination of the rest phyla. (C) Proteins in PF09828 are involved in the reduction of chromate accumulation and are essential for chromate resistance. Bacteria that hosts in plant produce the proteins identified as PF09828 to reduce the accumulation of chromate, resulting in the fast growth of the plant and preventing the transmission of cadmium to humans through the food chain leads to cadmium poisoning. For all the Pfam families which were foldable by C-I-TASSER, after aligning the Pfam species to the Interpro database, their protein functions were annotated by GO annotations, classified by three top annotation: Biological Process (D), Molecular Function (E), and Cellular Component (F).

To assess the utilization efficiency (*UE*) of metagenome sequences in Pfam structure modeling, we define *UE* = ∑*_i_*(*n_i_* /*N*). where *n_i_* is the number of sequences from the metagenome datasets that are homologous to the *i*th Pfam family, and *N* is the total number of metagenome sequences considered. In **Table 1** (**Column 7**), we list the *UE* values for different metagenome datasets on the Pfam families that are foldable by C-I-TASSER. It is shown that the utilization efficiencies of the three single biomes (Lake, Soil and Fermentor with *UE*=0.19, 0.49 and 0.94, respectively) are considerably higher than that from the combined dataset (0.15), although Gut’s *UE* are relatively low (0.04). If we count the number of Pfam families assisted by specific biomes, Soil and Fermentor assisted 907 and 2,000 families foldable per TB sequences, respectively, which are 2-5 times higher than that of the combined dataset, where the latter is comparable to those in previous metagenome structure modeling works (*10, 13*). These results suggest that targeted MSA collections from specific microbial biomes could improve the utilization efficiency of metagenome sequences compared to the approach that simply combines all available sequence datasets.

**Table 1.** Summary of utilization efficiency of metagenome sequences. For predicted Pfam families with unsolved structures, the statistic results for metagenome sequence utilization efficiency were calculated for results based on combined dataset (metagenomes from all of the four biomes), four single biomes, compared to datasets from previous studies.

| **Dataset** | **V<sup>a</sup>** | **Np<sup>b</sup>** | **Nf<sup>c</sup>** | **Nf/V<sup>d</sup>** | **Nf/Np<sup>e</sup>** | **UE<sup>f</sup>** | **P-value (IMG/Tera/Comb)<sup>g</sup>** |
| --- | --- | --- | --- | --- | --- | --- | --- |
| **IMG** | 1.41 | 2.25 | 614 | 435.5 | 272.9 | 0.10 | NA/NA/NA |
| **Tera Oceans** | 0.15 | 0.20 | 68 | 453.3 | 348.7 | 0.30 | NA/NA/NA |
| **Combined** | 2.4 | 4.3 | 1044 | 435.0 | 245.7 | 0.15 | 4E-19/E-26/NA |
| **Gut** | 1.4 | 2.1 | 105 | 75.1 | 50 | 0.04 | 5E-20/9E-18/6E-23 |
| **Lake** | 0.3 | 0.7 | 116 | 386.7 | 178.5 | 0.19 | 4E-22/3E-17/8E-22 |
| **Soil** | 0.68 | 1.4 | 617 | 907.4 | 440.7 | 0.49 | 9E-20/3E-19/5E-20 |
| **Fermentor** | 0.06 | 0.1 | 126 | 2000.0 | 1326.3 | 0.94 | 9E-21/5E-19/8E-20 |

V<sup>a</sup>: Volume size of protein sequence datasets (in TB)
Np<sup>b</sup>: Number of protein sequences (in billions)
Nf<sup>c</sup>: Number of foldable Pfam families with C-score >-2.5
Nf/V<sup>d</sup>: Number of foldable Pfam families/Volume size of dataset
Nf/M<sup>e</sup>: Number of foldable Pfam families/Number of sequences (per billion sequence)
UE<sup>f</sup>: Utilization efficiency of metagenome sequences in Pfam structure modeling (per 1000 metagenome sequences)
P-value<sup>g</sup>: P-value calculated on the UE values relative to 'IMG', 'Tera Oceans' and 'Combined data' respectively.

To decipher the important role of the solved Pfam families in their living environment, taxonomic profile and functional composition analyses were applied for each of the 964 Pfam families with single corresponding biome (**Figure 3**). The taxonomic profile for the 964 Pfam families illustrates specificity of contributions of microbial biome’s sequences to Pfam structure modeling (**Figure 3B**). Overall, similar to microbiome samples (**Figure 1B**), the heterogeneous species distribution reflects a biome-specific enrichment pattern for the 964 Pfam families. Moreover, the dominating species in specific Pfam families are often the dominating species in assisted microbiome samples for MSA constructions. For example, In Pfam families labeled with Gut biome (**Figure 1B** and **Figure 3B**), phylum Firmicutes and Bacteroidetes (both belonging to Gut) were the dominate phyla in Pfam families (0.41 ± 0.28 and 0.26 ± 0.14, respectively) and corresponding source biome (0.48±0.31 and 0.31±0.15, respectively), which indicates that this biome-specific enrichment pattern was influenced by the species composition of the microbiome samples.

In addition to structure modeling, the functional composition for the 964 Pfam families provides a useful insight into this biome-specific enrichment pattern. Based on the GO annotation, for example, 368 Pfam families were aligned to GO level-3 Biological Process (286), Molecular Function (90), and Cellular Component (189) (**Figure 3, D-F**). By analyzing the functional annotations for these Pfam families, the biome-specific enrichment pattern could also be detected, reflected by the fact that many function annotations were only detected in a single biome, including 129 (45.1%) for Biological Process, 69 (76.7%) for Molecular Function, and 109 (57.7%) for Cellular Component (**Table S3**).

Further functional analysis based on the biological process annotations reveals their important roles in helping the host species to adapt to their environment (**Figure 3D**). In Gut and Fermentor biomes, for example, the main functions are associated with Anaerobic energy metabolism (52.7% and 68.7% annotations for Gut and Fermentor, respectively). Enrichment of these functions could help their host to efficiently utilize the carbon sources to live in the oxygen-free environment and produce metabolites to interact with their host (*27, 28*). In the Lake biome, the main functions are associated with bacteria-specific cell motility (60.3%), to help their host adapt to the flowing water environment (*29*). Moreover, in the Soil biome, the functional roles of Pfam families with a C-score >= −2.5 were connected to such processes as nitrogen fixation (28.8%) and oxidation of iron (20.3%), sulfur (16.8%), and methane (10.3%), to take part in the soil chemical element cycle or adapt to the iron-enriched environment (*30–32*).

A typical example for supplementing homologous sequence for Pfam family 3D structure and function prediction is on a previously unsolved Pfam family PF09828, which contained 713 homologous sequences in the family and 98.3% sequences (701/731) of them are identified as bacteria (**Figure 3C**). After the sequences from the four biomes were included in the MSA construction search, the number of homologous sequences in the MSA for this family increased from 713 to 5,582 (Soil: 5,348, Lake: 151, Gut: 4, Fermentor: 79), resulting in a relatively high-accuracy contact-map and 3D structure prediction (C-score=-1.43). Interestingly, 526 sequences of the bacteria-sourced sequences in Pfam sequences (73.8%=526/713) are classified into phylum Proteobacteria, the dominant phyla in the Soil biome that counts for 93.0% of the homologous sequences supplemented in the MSA (**Figure 3B**). Further functional analysis reveals its role in the Soil biome: Bacteria that hosts in plant produce the proteins identified as PF09828 to reduce the accumulation of chromate in plant (*33*). The reduction of chromate in plant could promote the growth of plant and prevent the transmission of cadmium to humans through the food chain leads to cadmium poisoning (*34*). In **Text S1** and **Table S4**, we list ten other examples to showcase the biome-sequence-Pfam relationships. Taken together, these data illustrate potential correlations between the composition of the Pfam families and the source biomes used to supplement the MSAs for structure and functional modeling.

### Marginal effect analyses reveal biome-sequence-Pfam relationship

The results above have strongly indicated that the protein sequences from different biomes have profoundly different effects on supplementing homologous sequences of different Pfam families. To quantitatively examine the effect, we define the marginal effect of *i*th biome on *j*th Pfam family by *ME_ij_* = *n_ij_*/*m_j_*, where *n_ij_* is the number of homologous sequences for the *j*th family when searching the query through *i*th biome dataset, and *m_j_* is the number of homologous sequences in the *j*th family from the Pfam database. In **Table S5**, we list the marginal effects of the four biomes on all the 8,700 unknown Pfam families; the data shows that the contributions of different biomes to a specific Pfam can be drastically different, as reflected by their *ME* values. In **Figures 4A-D**, we present the contribution of biomes on the MSA collections for 4 examples from PF04213, PF10785, PF13864 and PF12357, where the microbiome samples were randomized for the MSA collections at different sequence numbers. For different Pfam families, the sequence homology pools are dominated by different biome datasets, suggesting again a strong link between biome and Pfam as regard to homologous sequence supplement.

**Figure 4.**
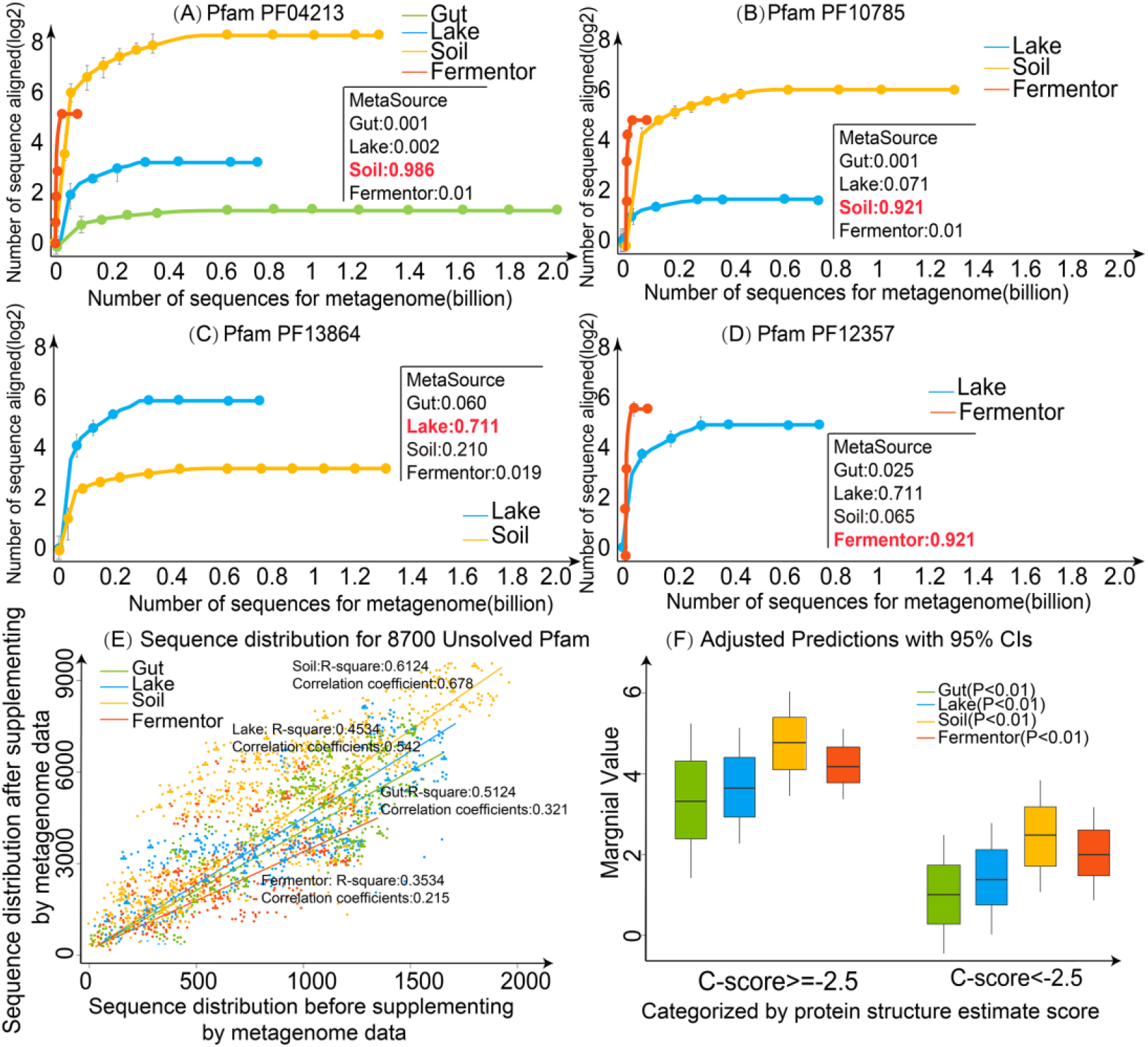
Evaluation of marginal effect for Pfam families. Collected from the four biomes, the homology sequences distribution of Pfam family (A) PF04213, (B) PF10785, (C) PF13864 and (D) PF12357 are illustrated, where the source biome of these Pfams was estimated by MetaSource. (E) The sequence distribution of metagenome data from the four biomes for all 8,700 Pfam families with unsolved structures. After the sequences from four biomes were aligned to 8,700 Pfam families with unsolved structures respectively, the marginal effect is estimated by comparison of the number of Pfam family’s homologous sequences before and after the use of the metagenome sequences. (F) marginal effect categorized by protein structure estimate scores.

To examine the overall trend of marginal effect, we plot in **Figure 4E** the (*n_ij_* + *m_j_*) vs *m_j_* values for all 8,700 Pfam families from 4 microbial biomes. For each biome, a linear regression was established based on the marginal value distribution of the 8,700 Pfam families. The correlation coefficients of simulated curve for four biomes are 0.678, 0.542, 0.321 and 0.215 for Soil, Lake, Gut and Fermentor, respectively, suggesting that the metagenome sequences are estimated to process a statistically positive effect to supplement the homologous sequences for Pfam families. On average, the marginal effect value is 5.28±3.25, 3.85±2.96, 3.48±3.11 and 4.12±1.65 for Soil, Lake, Gut and Fermentor, respectively. This rank of average marginal values for four biomes are largely consistent with the rank of species richness for the four biomes (**Figure S2**). Although the Soil biome has the highest overall marginal effect value, there are several hundreds of the Pfams families which have their highest marginal value from other three biomes, suggesting again the importance of biome-specific metagenome sequence selection to maximize the efficiency of MSA collection.

In **Figure 4F**, we split Pfam families into two groups based on C-I-TASSER folding results. It was shown that the *ME* value for the families with C-score ≥-2.5 is much higher than that with C-score<-2.5 (5.27±3.44 vs. 1.28±0.85 with a p-value=3.86e-26 in Student’s t-test). Therefore, marginal effect value is also strongly correlated with the ability of biome-specific metagenome sequence to assist the 3D structure assembly simulation through supplementing more homologous sequences.

### MetaSource prediction model for effective homologous sequence supplements

A targeted model with correct metagenome selection could effectively assist the homologous sequences supplement and 3D structure prediction for unsolved Pfam families. Here we proposed the MetaSource prediction model to identify one or a set of biomes which can better supplement homologous sequences for specific Pfam families. First, to determine whether the source biome of the query Pfam family is one of the four biomes, a binary classification model was constructed, by using the 964 Pfam families labeled with a single biome as the training dataset, and 7,736 (=8,700-964) Pfam families with unsolved structures as the testing dataset. As shown in **Figure 5A**, MetaSource achieves an AUC of 0.96 under 0.001 permutation p-value on the binary classification test. Second, to predict the most probable source biome out of the four biomes for a Pfam family, the multi-class Random Forest algorithm was chosen to construct this model. In this context, biome that could supplement the largest number of homologous sequences were considered as “correct” biome. The 964 Pfam families labeled with a single biome was used, with 20 cross-validation iterations (**Figure 5B**), showing a strong predictive power of MetaSource for the Pfam families, with a micro-average AUC of 0.94, under 0.001 permutation p-value. The top 20 important features used in MetaSource were supplied in **Figure S3**.

**Figure 5.**
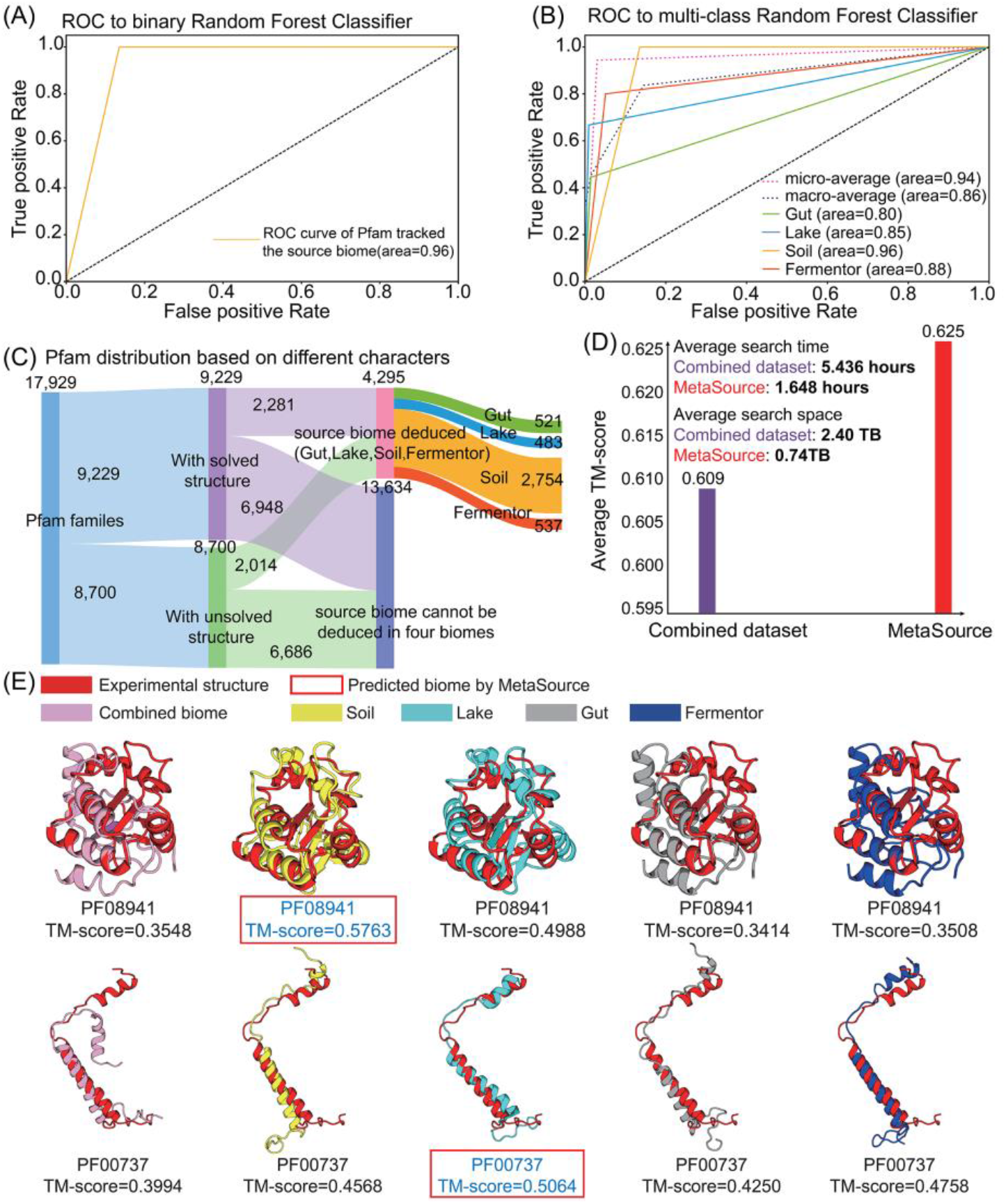
The source biomes predicted by MetaSource for Pfam families. (A) The ROC analysis of binary-classification MetaSource model. This model was constructed to determine whether the source biome of the query Pfam family is one of the four biomes. (B) The ROC analysis of multiple-classification MetaSource model. This model was constructed to predict the source biome for Pfam families. To evaluate the overall prediction accuracy, the micro-average (obtained by aggregating the contributions of all classes to compute the average metric) and macro-average value (calculated by the metric independently for each class and take the average) were applied. (C) The Pfam classification result for all the Pfam families based on the prediction result of MetaSoure model. (D) Average TM-score and average MSA search time for the combined and MetaSource predicted biome datasets. (E) Case studies of modeling Pfam (PF08941 and PF00737) with MSA from different biomes. The model with the highest TM-score is shown in blue font. The model labeled with red frame is the source biome predicted by MetaSource.

To further examine the practical usefulness of the MetaSource model in 3D structure modeling, we incorporated the Pfam families with known structure into our analysis as the validation dataset **(Figure 5C)**. First of all, as listed in **Table S6**, MetaSource was able to predict the biome which is identified with highest *Neff* (or based on TM-score) with an accuracy of 79.9% (or 80.2% for TM-score) (permutation p-value: 0.001). In **Figure 5D**, we compare the average TM-score of the C-I-TASSER models when using MSAs collected from the combined dataset and the dataset chosen by MetaSource. It was shown that, although the volume of the sequence database is much smaller (0.74 TB/per target and 2.40 TB/per target for MetaSource and Combined datasets respectively), using the targeted dataset from MetaSource results in a higher TM-score (0.625) than that of the combined dataset (0.609), which corresponds to a p-value=6.3E-6 in Student’s t-test. Accordingly, the speed of MSA search based on MetaSource (1.65 hours/per target) is also faster than that from the combined dataset (5.44 hours/per target). The result may be understandable because sequences from the “wrong” source biome can produce “noise” to the MSA collection and deep-learning-based contact prediction, where the identification of correct source biomes could help depress such noises.

In **Figure 5E**, we present two Pfam examples from PF08941 and PF00737 with known structure, for which MetaSource predicted Soil and Lake as the best source biome, respectively. In both cases, only the models with the MetaSource predicted biomes could create a model with TM-score above 0.5. We also noticed that, although the MSA from the combined biome contains more sequences than single biome, the structure models are clearly worse than the MSA from some single biome (Soil or Lake), probably due to the noise contribution from irrelevant metagenome sequences. Taxonomic profile analyses also showed that PF08941 and PF00737 are mainly composed with proteins from phylum Proteobacteria and Cyanobacteria, which dominate in Soil and Lake biomes, respectively(*22, 35*). Taken together, these results demonstrated again that the intrinsic links among biome-sequence-Pfam can be used as a useful means to enhance effectiveness and efficiency of the targeted approach for metagenome data on protein structure prediction and function annotations.

## Conclusion

As a grand reservoir of novel genes and proteins, microbial communities contain a large number of uncultured species that are unique for adapting their living environments. Nowadays, the metagenome sequencing technology has been advanced enough to sequence microbial communities in many of the known biomes on Earth, while more complete gene catalogs of microbial communities have been obtained from some biomes than others due to the accessibility of the species as well as their functional genes. While these microbiome sequences have been shown useful for boosting the accuracy and capacity of deep learning-based protein structure and function predictions, the model training and metagenome search were largely blind and fall short in efficiency in source tracking the most relevant biome datasets for specific protein targets. For designing a more effective targeted approach, deeper insights should be obtained to link microbiome biomes with protein family homologous sequences.

In this study, we utilized 2.4 TB of the microbiome sequencing data, representing 4.25 billion microbiome sequences, to investigate the usefulness of metagenome sequences from specific biomes for protein structure prediction of individual Pfam families. The microbiome sequences from the four biomes (Gut, Lake, Soil and Fermentor) boosted multiple sequence alignments with credible multiplicity for 2,214 out of 8,700 Pfam families with unsolved structures. By applying C-I-TASSER *ab initio* structure folding pipeline, highly reliable folds were constructed for 1,044 Hard Pfam families, which account for 12% of all unknown Pfam families.

To further examine the association between the metagenome sequences and Pfam families, we quantified the marginal effect of metagenome sequences on Pfam families, where the data shows that metagenome sequences from different biomes have drastically discriminable power to different Pfam families. Accordingly, a machine-learning model, MetaSource, was constructed for source tracking the most relevant biome datasets for specific Pfam family structure modeling. The utilization of the MetaSource predicted biomes have resulted in 3.2-fold reduction in the database size and 3.3-fold increase in MSA construction speed, but with 3% of TM-score increase in the C-I-TASSER final models. This result is particularly encouraging in this postgenomic era when the number of genome and metagenome sequences increases exponentially, and the speed and memory requests become a major bottleneck for sequence mining and MSA collection through large-scale sequence database searching (*8*). These findings could be used as a useful bluebook to guide the modeling of protein structure and function based on the deeper insights into the biome-protein association.

Finally, we should emphasize that this study only considers four microbiome biomes (Gut, Lake, Soil and Fermentor) with C-I-TASSER structure modeling method as an illustration. Much more metagenome datasets could be straightforwardly incorporated into this model. Moreover, with the rapid progress of the field, C-I-TASSER considering only contact-map restraints no longer represents the state of the art of protein structure prediction. With more advanced methods, including Alphafold (*36*), D-I-TASSER (*37*) and trRosetta (*38*), which consider more thorough sequence-based restraints armed with more advanced deep-learning methods, we expect that the targeted metagenome selection approach should have more sensitive and pronounced impact on the efficiency and effectiveness of the protein structure prediction and structure-based protein function annotations.

## Author contributions

K.N. and Y.Z. conceived the project and designed the experiment; P.Y and W.Z. developed methods and perform experiments; All authors wrote the manuscript and approved the final submission.

## Data and materials availability

All data is available in the main text or the supplementary materials.

## References

1. D. Baker, A. Sali, Protein structure prediction and structural genomics. Science 294, 93–96 (2001).

2. Y. Zhang, Progress and challenges in protein structure prediction. Curr Opin Struct Biol 18, 342–348 (2008).

3. K. T. Simons, C. Kooperberg, E. Huang, D. Baker, Assembly of protein tertiary structures from fragments with similar local sequences using simulated annealing and Bayesian scoring functions. J Mol Biol 268, 209–225 (1997).

4. D. Xu, Y. Zhang, Ab initio protein structure assembly using continuous structure fragments and optimized knowledge-based force field. Proteins 80, 1715–1735 (2012).

5. J. Yang et al. The I-TASSER Suite: protein structure and function prediction. Nat Methods 12, 7–8 (2015).

6. S. Wang, S. Sun, Z. Li, R. Zhang, J. Xu, Accurate De Novo Prediction of Protein Contact Map by Ultra-Deep Learning Model. PLoS Comput Biol 13, e1005324 (2017).

7. Y. Li et al., Deducing high-accuracy protein contact-maps from a triplet of coevolutionary matrices through deep residual convolutional networks. PLoS Comput Biol, doi: https://doi.org/10.1371/journal.pcbi.1008865 (2021).

8. C. Zhang, W. Zheng, S. M. Mortuza, Y. Li, Y. Zhang, DeepMSA: constructing deep multiple sequence alignment to improve contact prediction and fold-recognition for distant-homology proteins. Bioinformatics 36, 2105–2112 (2020).

9. R. Shrestha et al., Assessing the accuracy of contact predictions in CASP13, (ranking result can be seen at: http://www.predictioncenter.org/casp13/zscores_rrc.cgi). Proteins 87, 1058–1068 (2019).

10. S. Ovchinnikov et al., Protein structure determination using metagenome sequence data. Science 355, 294–298 (2017).

11. C. UniProt, UniProt: a worldwide hub of protein knowledge. Nucleic Acids Res 47, D506–D515 (2019).

12. A. T. Brunger et al., Crystallography & NMR system: A new software suite for macromolecular structure determination. Acta Crystallogr D Biol Crystallogr 54, 905–921 (1998).

13. Y. Wang et al., Fueling ab initio folding with marine metagenomics enables structure and function predictions of new protein families. Genome Biol 20, 229 (2019).

14. I. A. Chen et al., The IMG/M data management and analysis system v.6.0: new tools and advanced capabilities. Nucleic Acids Res, (2020).

15. E. W. Sayers et al., GenBank. Nucleic Acids Res 48, D84–D86 (2020).

16. L. A. Abriata, G. E. Tamo, M. Dal Peraro, A further leap of improvement in tertiary structure prediction in CASP13 prompts new routes for future assessments. Proteins-Structure Function and Bioinformatics 87, 1100–1112 (2019).

17. W. Zheng et al., Deep-learning contact-map guided protein structure prediction in CASP13. Proteins 87, 1149–1164 (2019).

18. A. L. Mitchell et al., MGnify: the microbiome analysis resource in 2020. Nucleic Acids Res 48, D570–D578 (2020).

19. D. Ramanan et al., Helminth infection promotes colonization resistance via type 2 immunity. Science 352, 608–612 (2016).

20. P. B. Eckburg et al. Diversity of the human intestinal microbial flora. Science 308, 1635–1638 (2005).

21. G. P. Donaldson et al., Gut microbiota utilize immunoglobulin A for mucosal colonization. Science 360, 795–800 (2018).

22. P. Poole, V. Ramachandran, J. Terpolilli, Rhizobia: from saprophytes to endosymbionts. Nat Rev Microbiol 16, 291–303 (2018).

23. A. Detman et al., Cell factories converting lactate and acetate to butyrate: Clostridium butyricum and microbial communities from dark fermentation bioreactors. Microb Cell Fact 18, 36 (2019).

24. W. Zheng et al., LOMETS2: improved meta-threading server for fold-recognition and structure-based function annotation for distant-homology proteins. Nucleic Acids Res 47, W429–W436 (2019).

25. Y. Zhang, J. Skolnick, TM-align: a protein structure alignment algorithm based on the TM-score. Nucleic Acids Res 33, 2302–2309 (2005).

26. H. C. Flemming, S. Wuertz, Bacteria and archaea on Earth and their abundance in biofilms. Nat Rev Microbiol 17, 247–260 (2019).

27. K. Fenn et al., Quinones are growth factors for the human gut microbiota. Microbiome 5, 161 (2017).

28. T. Ito et al., Genetic and Biochemical Analysis of Anaerobic Respiration in Bacteroides fragilis and Its Importance In Vivo. mBio 11, (2020).

29. K. L. Hentchel et al., Genome-scale fitness profile of Caulobacter crescentus grown in natural freshwater. ISME J 13, 523–536 (2019).

30. K. A. Weber, L. A. Achenbach, J. D. Coates, Microorganisms pumping iron: anaerobic microbial iron oxidation and reduction. Nat Rev Microbiol 4, 752–764 (2006).

31. N. Fierer, Embracing the unknown: disentangling the complexities of the soil microbiome. Nat Rev Microbiol 15, 579–590 (2017).

32. J. K. Jansson, K. S. Hofmockel, Soil microbiomes and climate change. Nat Rev Microbiol 18, 35–46 (2020).

33. G. Sturm et al., Chromate Resistance Mechanisms in Leucobacter chromiiresistens. Appl Environ Microbiol 84, (2018).

34. Y. Wang et al., Characteristics and in situ remediation effects of heavy metal immobilizing bacteria on cadmium and nickel co-contaminated soil. Ecotoxicol Environ Saf 192, 110294 (2020).

35. P. J. Cabello-Yeves, F. Rodriguez-Valera, Marine-freshwater prokaryotic transitions require extensive changes in the predicted proteome. Microbiome 7, 117 (2019).

36. A. W. Senior et al., Improved protein structure prediction using potentials from deep learning. Nature 577, 706–710 (2020).

37. Y. Li et al., Protein 3D Structure Prediction by Zhang Human Group in CASP14. Abstract of 14th Critical Assessment of Structure Prediction, 328 (2020).

38. J. Yang et al., Improved protein structure prediction using predicted interresidue orientations. Proceedings of the National Academy of Sciences of the United States of America 117, 1496–1503 (2020).

