## Supplementary material for "Decoding microbiome and protein family linkage to improve protein structure prediction": Materials and Methods, Supplementary Texts, Supplementary Figures, Supplementary Tables

### Supplementary Materials

#### Table of Content

**Section 1: Materials and Methods**

1.1 Microbial community cohorts collected from four biomes

1.2 Taxonomic and functional analysis for Pfam families

1.3 Pfam family dataset construction for 3D structure modeling

1.4 Procedures of the multiple sequence alignment collection

1.5 Contact-assisted structure prediction by C-I-TASSER

1.6 C-I-TASSER model quality estimation

1.7 MetaSource model construction and evaluation for predicting the source biome of Pfam families

**Supplementary Texts**

**Text S1**. Case studies verified the applicability and interpretability of the targeted MetaSource model

**Text S2**. The normalized number of effective sequences (*Neff*) in MSA

**Supplementary Figures S1-S9**

**Supplementary Tables S1-S6**

**References**

#### Section 1: Materials and Methods

##### 1.1 Microbial community cohorts collected from four biomes

We collected metagenome data from the EBI database (https://www.ebi.ac.uk/metagenomics/). Referred to the EBI database, the microbial niches were annotated in a hierarchical classification tree, named as biome (1). Hence, to cover all the typical biomes, samples under three top-layer biomes were screened: "Engineered" biome (the affiliate biome "Fermentor" was selected as a representative biome), "Environmental" biome (the affiliate biome "Soil" and "Lake" were selected as representative biomes), and "Host-associated" biome ("Gut" biome as representative biome). The samples from Gut biome were collected from human gut covering different countries (**Figure S4**) and animal (mice, pigs, cattle, etc.) intestines.

Since the EBI data has been processed by different processing pipeline, we reanalyzed the 1,705 metagenomes uniformly using pipeline version 4.1. If the data are processed by the pipeline older than version 4.1, the raw reads were downloaded and performed by the SeqPrep (version 1.2) and Trimmomatic (version 0.35) for quality control. The proteins were then predicted by FragGeneScan (version 1.20) and Prodigal (version 2.6.3). Finally, a total of 4.25 billion protein sequences were collected from the 1,705 high-quality samples. Moreover, the taxonomic profiles were predicted by MAPseq (version 1.2.2) and the functional composition was calculated by InterProScan based on GO annotation (version 5.25-64.0).

##### 1.2 Taxonomic and functional analysis for Pfam families

To decipher the association between microbial communities and Pfam families, the taxonomic and functional distributions for the Pfam families was analyzed. Since the original species for every homology sequence of a Pfam family could be tracked, all the species information (species classification and count number for each species) was obtained from the Pfam database and used as the taxonomic profile. Moreover, the InterPro annotation and their associated GO terms for each family were used as the functional annotation which was stored in Pfam database. All these data are available at the FTP site, under the release version 32.0 (ftp://ftp.ebi.ac.uk/pub/databases/Pfam/releases/Pfam32.0/).

##### 1.3 Pfam family dataset construction for 3D structure modeling

Pfam (version 32) is a database that contains 17,929 protein families, each represented by a hidden Markov model (HMM). Typically, each Pfam entry is comprised of a seed alignment, which forms the basis to build a profile hidden Markov model using HMMER (2). The profile HMM is then queried against a sequence database, and all matches scoring above the curated threshold are aligned back to the profile HMM to generate the full alignment. Pfam includes 9,229 protein families that have at least one member with experimentally determined structures. For the remaining 8,700 families, there is no structural information available for any member, where a breakdown of the Pfam families in this study is shown in **Figure S5**.

372 Pfam families has been randomly selected from the 9,229 known families as benchmark dataset to investigate the performance of C-I-TASSER and MetaSource. The families are Hard targets as defined by LOMETS(3) since there is no homologous template with a sequence identity <30% in the PDB library. To generate a representative sequence, we search each of the Pfam families against the SCOPe database, where the best hit with the PDB ID appearing in the Pfam structure member list will be selected. Finally, 168 Pfam families are used for benchmarking C-I-TASSER

and the remaining 204 families for testing MetaSource.

Out of the 8,700 unknown families, we first removed the entries with less than 50 amino acids of sequence length, resulting in a set of 8,266 Pfam families. We select one representative sequence for C-I-TASSER modeling for each given Pfam family. To do so, we first ran 'HMMsearch' to search the family against the UniRef100 database, with the sequences hit ranked by their E-values. For the best hit with the lowest E-value, we run DeepMSA (4) to build MSA, where 2,251 Pfam families with *Neff* score >16 (see *Neff* definition in **Eq. S3** in **Text S2**) and defined as "Hard" targets by LOMETS are selected for modeling. Finally, the 1,044 Pfam families with model having C-score ≥-2.5 from C-I-TASSER are selected for training MetaSource model (**Figure S5**).

##### 1.4 Procedures of the multiple sequence alignment collection

To predict the structure and function of the 8,700 unknown Pfam families, the metagenome database of a combination of the four biomes (Gut, Lake, Soil and Fermentor) were attempted to supplement the Pfam homologous sequence using a 3-step procedure outline by DeepMSA (4) (**Figure S6**). In step 1, HHblits (5) from HH-suite is used to search the query sequence against UniClust30 (6) to generate the first-level MSA. In step 2, the Jackhmmer from the HMMER (2) package is used to search the query sequence against UniRef90 (7) to extract full-length sequences (hits) and HHblits is used to convert the full-length sequences into a custom HHblits format database. Starting from the first level MSA, HHblits is again applied to search this custom database to generate the second level MSA. In step 3, the second level MSA is converted by hmmbuild from the HMMER package into a Hidden Markov Model (HMM) and the HMM is then searched against the metagenome sequence database (the combination of the four biomes) by HMMsearch from the HMMER package to extract full-length hits. Similar to step 2, hits from HMMsearch are built into a custom HHblits database. The second level MSA is used to jump-start an HHblits search against this new custom HHblits database to get the third level MSA. For each MSA, a *Neff* score is computed by **Eq. S3,** where the families with *Neff* score ≥16 are selected as 'effective Pfam families'. Finally, the homologous sequences in the MSA are collected for further protein structure modeling.

The same MSA construction pipeline is also used for MSA generation for 168 C-I-TASSER benchmark dataset and 204 MetaSource testing dataset. Additionally, for MetaSource testing dataset, we also generated four sets of MSAs which used four biomes as database individually instead of the combined metagenome in DeepMSA step 3. Those five sets MSAs (from four biomes individually and the combined one) of MetaSource testing dataset are used for building five sets of C-I-TASSER models to check the correctness of MetaSource.

##### 1.5 Contact-assisted structure prediction by C-I-TASSER

Based on the collected MSAs, residue-residue contact-maps are predicted using 5 deep-learning and co-evolution based predictors, including TripletRes (8), ResTriplet (9), NeBcon (10), ResPRE (11), and ResPLM (12). The consensus contacts are collected from top $L$ contacts from the five predictors, respectively. These contacts are implemented in the C-I-TASSER simulation through the following potential:

$$E_{contact}(d_{ij}) = \begin{cases} -U_{ij}, & d_{ij} \le 8 \\ -\frac{U_{ij}}{2} \cdot \left[1 - \sin\left(\frac{d_{ij} - (8 + D)/2}{D - 8} \cdot \pi\right)\right], & 8 < d_{ij} < D \\ \frac{U_{ij}}{2} \cdot \left[1 + \sin\left(\frac{d_{ij} - (80 + D)/2}{80 - D} \cdot \pi\right)\right], & D < d_{ij} < 80 \\ U_{ij}, & d_{ij} \ge 80 \end{cases} \quad \text{(S1)}$$

where $d_{ij}$ is the $C\beta$ distance between residue pair *i* and *j*; $U_{ij}$ is the contact prediction confidence score for this residue pair; *D* is a protein length-dependent parameter to change the gradient of the well which ranges from 14 to 18 Å (**Figure S7**).

Starting from the representative sequence of a Pfam family, homologous templates are detected from the PDB library by LOMETS meta-threading server (3), which consisting 11 individual threading programs, CEthreader (13), CNFsearch (14), FFAS3D (15), HHpred (16), HHsearch (16), MUSTER (17), Neff-MUSTER (18), PPAS (19), PROSPECT2 (20), SP3 (21), and SparksX (22). The consensus contact and distance restraints are collected from the LOMETS template alignments, which are combined with the sequence-based contact potential as **Eq. S1** to guide the I-TASSER structural assembly simulations (18). Finally, the decoy conformations from the C-I-TASSER simulation trajectories are clustered by SPICKER (23), where the largest cluster is further refined at atomic-level by FG-MD (24) and returned as the final model.

##### 1.6 C-I-TASSER model quality estimation

To estimate the model quality, we run C-I-TASSER on 168 Pfam proteins that have known structures in PDB, where all 168 proteins were "Hard" targets according to LOMETS classification (19). The data in **Figure S8** demonstrate a strong correlation between the TM-score of the C-I-TASSER models and the confidence score (C-score) of the folding simulations, which has a Pearson correlation coefficient (PCC=0.801). Here, the confidence score is defined by

$$\text{C-score} = w_1 * \ln\left(\frac{1}{\text{K}}\sum_{\text{i}=1}^{\text{K}} \frac{Z(i)}{Z_0(i)}\right) + w_2 * \ln(Sr) + w_3 * \ln(Dc) \quad (S2)$$

where $Z(i)$ and $Z_0(i)$ are the highest Z-score of the templates by the $i$-th LOMETS2 threading program and the corresponding Z-score cutoff for distinguishing between good and bad templates. These Z-score related parameters describe the significance of the LOMETS threading alignments. $Sr$ is the satisfaction rate of top-$L$ long-range contacts in the final model, i.e., $Sr = 1/n_L \sum_{i=1}^{n_L} \delta_i$, where $n_L$ is the number of the top-$L$ predicted contacts with residue separation >24, $\delta_i = 1$ (or 0) if the $i$-th contact is satisfied (or not satisfied) in the final C-I-TASSER model. $Dc$ measures the degree of structure convergence in the C-I-TASSER simulation and is calculated by $Dc = \frac{M}{M_{tot}}/\langle R\rangle$, where $M$ is the number of decoys in the SPICKER cluster, $M_{tot}$ is the total number of structure decoys generated in C-I-TASSER simulation, and $\langle R\rangle$ is the average RMSD of the structure decoys to the cluster centroid. Weight parameters ($w_1 = 1.36$, $w_2 = 0.67$, $w_3 = 0.77$) are decided by maximizing the PCC. If we select a C-score cutoff of -2.5, the Matthews correlation coefficient (MCC) on the benchmark dataset reached a maximum of 0.614 and an FDR of only 6.96% (**Figure S8**).

##### 1.7 MetaSource model construction and evaluation for predicting the source biome of Pfam families

To identify the source biome that has the largest number of homologous sequences for a given Pfam family, we construct a machine learning model named MetaSource (https://github.com/HUST-NingKang-Lab/MetaSource). As depicted in **Figure S9**, the pipeline consists of 4 consecutive steps:

1. Investigation of the biome-sequence-Pfam association: The sequences collected from four biomes (Gut, Lake, Soil and Fermentor) were used for supplementing the homologous sequences of Pfam families (**Figure S9A**). Furthermore, by comparing the homologous sequence number before and after supplementing the metagenome sequence for Pfam families, the marginal effect analysis was applied to perform a quantitative assessment on the biome-sequence-Pfam association (**Figure S9C**).

2. Training dataset construction using Pfam families with unsolved structure (**Figure S9D**): The Pfam families foldable by C-I-TASSER (e.g., C-score ≥-2.5) are used as the training dataset for the prediction model since the biome genomes have a stronger contribution to the MSA construction for these Pfam families (**Figure S9B**). For the Pfam families foldable by C-I-TASSER, the biome with highest *Neff* was used as the data label after supplementing the homology sequences from four biomes respectively. And the taxonomic profile on genus level for Pfam families were used as the features for training set. To reduce the complexity of the data, the genera with an average relative abundance less than 0.001 were filtered out. Furthermore, to select the features with a significant difference in the distribution of multiple groups, the Kruscal-Wilcox test was performed with a p-value over 0.05 by and q-value over 0.05 calculated by the Bonferroni method.

3. Constructing the MetaSource model (**Figure S9E**): Given our relatively small dataset, we sought to identify a model that would tend toward low variance and the Random-Forest algorithm was applied. Firstly, to predict whether the source biome could be one of the four common biomes (Gut, Lake, Soil, Fermentor), a binary Random-Forest algorithm was applied. The positive dataset was 964 Pfam families foldable by C-I-TASSER and the negative dataset was 7,736 (=8,700-964) Pfam families with unsolved data. Then, to predict the single biome that could effectively supplement homologous sequences for the specific Pfam family, a multi-label Random-Forest classifier was applied using 964 Pfam families foldable by C-I-TASSER as training data. To find the best combination of model parameters, grid search was applied to exhaustively search over all parameter values. Then, the model was trained by 20 cross-validation iterations, and in each iteration, the model was trained on three-fourths of the dataset. Finally, the capacity to predict the source biome that was left out was assessed.

4. Validating the MetaSource using the Pfam families with solved structure (**Figure S9F**): To validate the performance of MetaSource prediction model, the Pfam families with solved structure was used to evaluate the performance of MetaSource on prediction of source biome. For each Pfam family with known structure, we built MSA after querying the homology sequences from Gut, Lake, Soil, Fermenter and combined four biomes, respectively. Those five MSAs were passed to C-I-TASSER to build five structures individually and then were compared with experimental structure. If the biome with highest *Neff* is consistent with the prediction source biome using MetaSource, the prediction will be considered correct. All the four steps are implemented by Python, using the scikit-learn package (version 0.22, https://scikit-learn.org/stable/).

#### Supplementary Texts

##### Text S1. Case studies verified the applicability and interpretability of the targeted MetaSource model

Through the case studies, our targeted metaSource model shows a strong applicability and good biological interpretability. Among 964 Pfam families (*Neff* over 16 and C-score over -0.25), 10 Pfam families are selected for case studies (**Table S4**).

These Pfam families are selected based on the literature review and the comparison of prediction result (measured by the *Neff* score) for four commonly used datasets (Uniref100(25), IMG(25), Tara Oceans (26) and Metaclust (26)). These four datasets are commonly used to assist the structure and function prediction for unsolved proteins.

First, assisted by Soil biome, PF05120 could be supplemented with sufficient homologous sequences (*Neff* score=487.5 and C-score=-0.18). Based on our targeted metasource model, this Pfam family is successfully classified into the Soil biome (accuracy:0.968). However, the other four commonly used datasets supply insufficient homologous sequences, reflected by the lower *Neff* score than the Soil biome used in our research: 32.0, 336.8, 69.0 and 178.9 for Uniref100, IMG+ Uniref100, Tara Oceans+Uniref100 and Metaclust+Uniref100, respectively. This result indicates that our targeted prediction model can accurately predict that Soil biome could be used to supplement the homologous sequence of PF05120. Furthermore, this prediction result could be interpreted by its unique biological role in Soil biome for PF05120: According to the records in Pfam families, the members in PF05120 are annotated as gas vesicles proteins. These gas vesicles proteins are permeable to ambient gases by diffusion and provide buoyancy, enabling cells to move based on the air-soil interface(27,28). This protein plays an important role in the communications between different soil microbiome communities(27).

Second, the accuracy and interpretability of our targeted metaSource model could also be proved by other biomes: Among the 964 Pfam families with C-score >-0.25, PF12652 is successfully classified into the biome of Fermentor by our metasource model (accuracy:0.975). Actually, measured by a high *Neff* score (305.6) and C-score (-0.16), our protein structure prediction results also confirm this result. However, insufficient homologous could be provided by four datasets for PF12652 (Nf score 232.6,264,295.2,299.6 for Uniref100, IMG+ Uniref100, Tara Oceans+ Uniref100 and Metaclust+ Uniref100, respectively). The fermentor-related function of proteins in PF12652 could explain this result: based on the records in PF12652, this Pfam family is related to spore development. The bacteria that enrich the spore development function are closely related to anaerobic fermentation, the main function of fermenters (29).

Finally, based on an investigation of the correctly classified Pfam families, great application prospects have sprung up using our targeted MetaSource model: PF13822(classified into Soil biome, accuracy:0.982) is identified as Acyl-CoA carboxylase epsilon subunit, which is involved in the biosynthesis of long-chain fatty acids. The long-chain fatty acids are important for Rhizobium leguminosarum Growth and Stress Adaptation(30). PF09828 and PF05425 are two important antibiotics. These two antibiotics shows the resistance to chromate and copper, which are harmful to the agricultural plants and human (31,32). PF09650(classified into Soil biome, accuracy:0.965), is identified as putative polyhydroxyalkanoic acid (PHA) system protein, and could produce the bioplastic(33).

**Text S2. The normalized number of effective sequences (*Neff*) in MSA**

The depth of a multiple sequence alignment (MSA) is measured by the normalized number of effective sequence (*Neff*) in this work:

$$Neff = \frac{1}{\sqrt{L}} \sum_{i=1}^{N} \frac{1}{1 + \sum_{j=1, j \neq i}^{N} I\left[S_{j,i} \geq 0.8\right]} \qquad (S3)$$

where $L$ is the length of protein, $N$ is the number of sequences in the MSA, $S_{j,i}$ is the sequence identity between the *j*-th and *i*-th sequences. $I\left[S_{j,i} \geq 0.8\right]$ equals to 1 if $S_{j,i} \geq 0.8$, or zero otherwise. Therefore, *Neff* is essentially equal to the number of non-redundant sequences (sequence identity<0.8) in the MSA normalized by the protein length.

#### Supplementary Figures

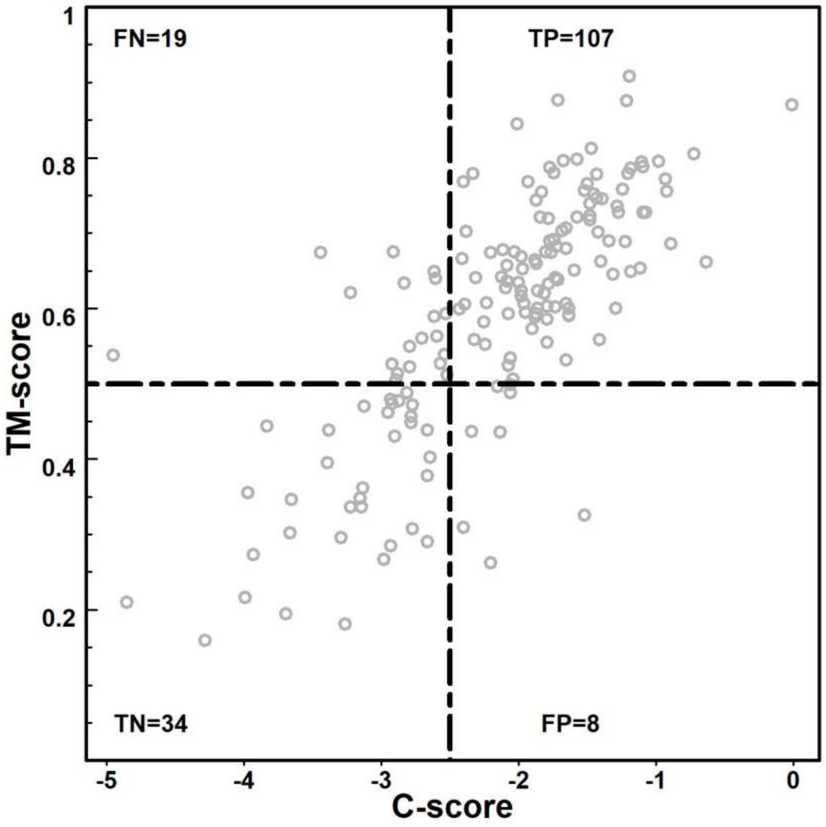

**Figure S1. Accuracy estimation of predicted models using C-score defined by Eq. S2**, represented by TM-score of the first C-I-TASSER model versus C-score.

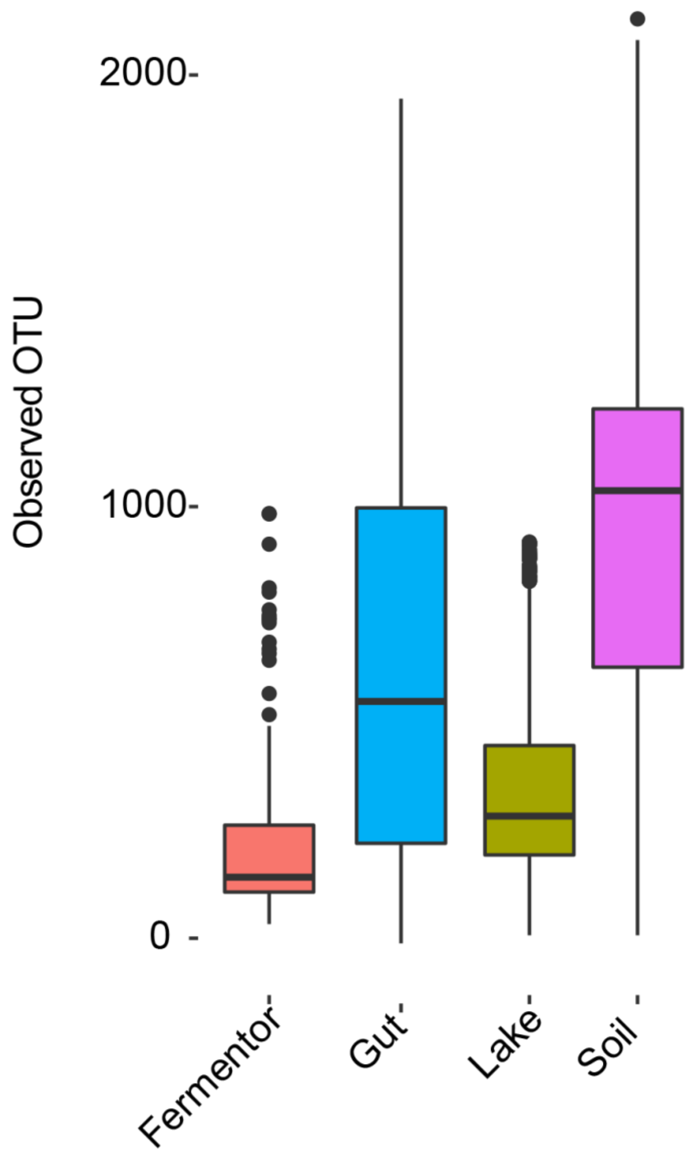

**Figure S2. The species richness statistic for four biomes (Fermentor, Gut, Lake and Soil).** The raw metagenome sequences were assembled, extract the 16s rRNA and clustered by 97% similarity to obtain the operational taxonomic units (OTUs) distribution, sequentially. The OTU distribution could represent the species richness in corresponding environment.

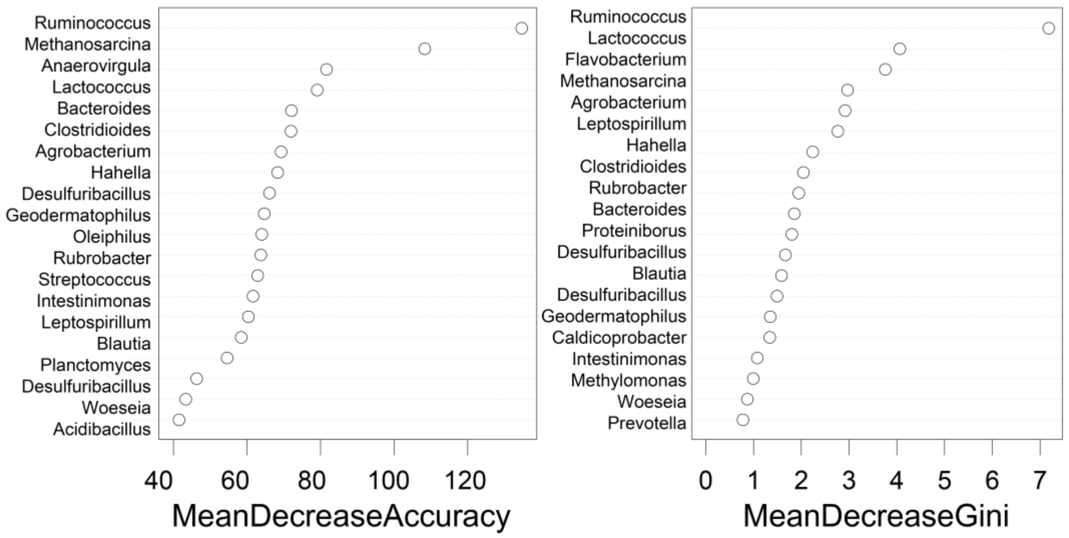

**Figure S3. The top 20 importance features (genus) for the multiple-classified Random Forest model**. The importance of features was estimated and ranked by accuracy and Gini index.

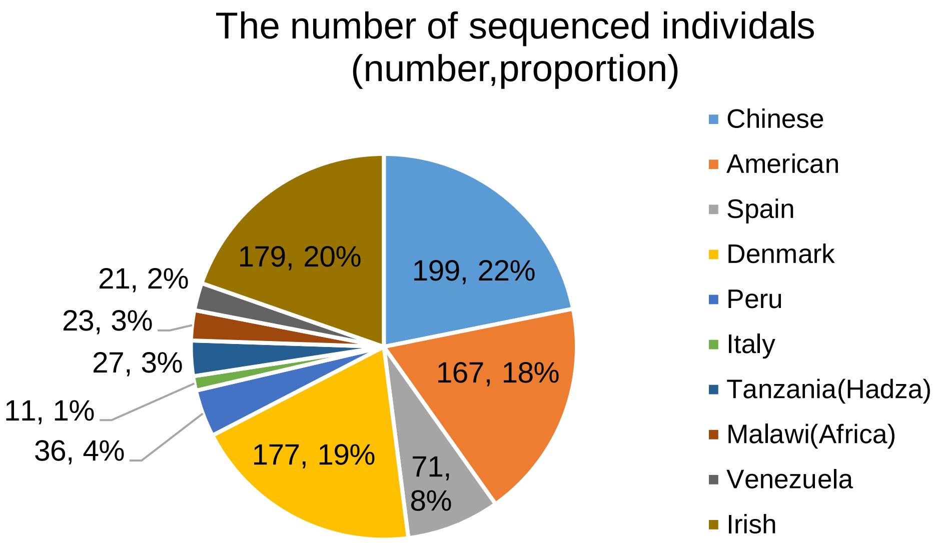

**Figure S4. For the Gut biome, the statistical result based on country distribution.** The 911 samples were collected from 10 countries, covering four continents (Africa, Asia, Europe, Americas).

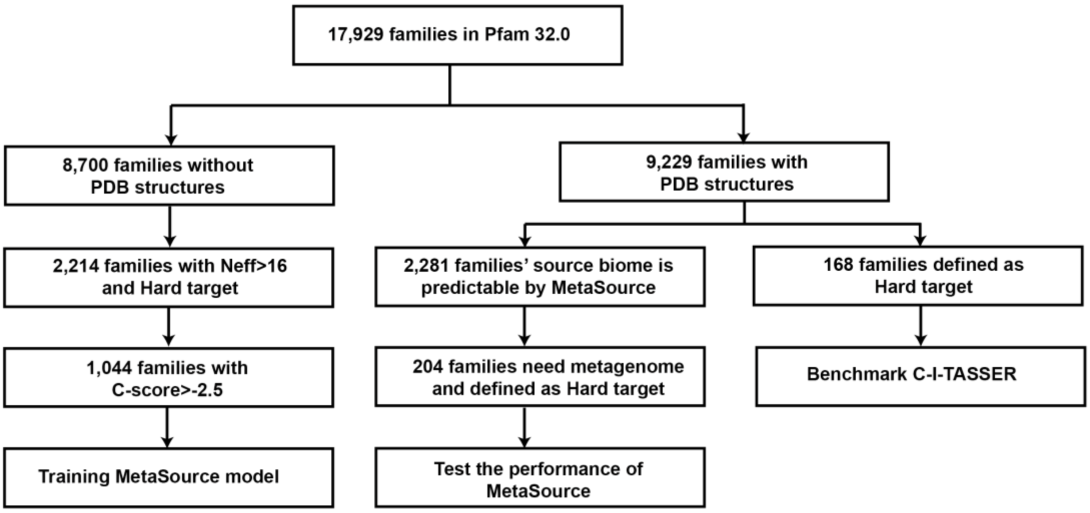

**Figure S5. Data collection flow from Pfam database for training and testing MetaSource and benchmarking the C-I-TASSER.** For 8,700 Pfam families with unsolved structure, 1,044 Pfam families were used to train the MetaSource prediction model after a set of filtration. For 9,229 Pfam families with solved structure has been randomly selected as benchmark dataset to investigate the fold ability of C-I-TASSER, and testing dataset for qualify the performance of MetaSource.

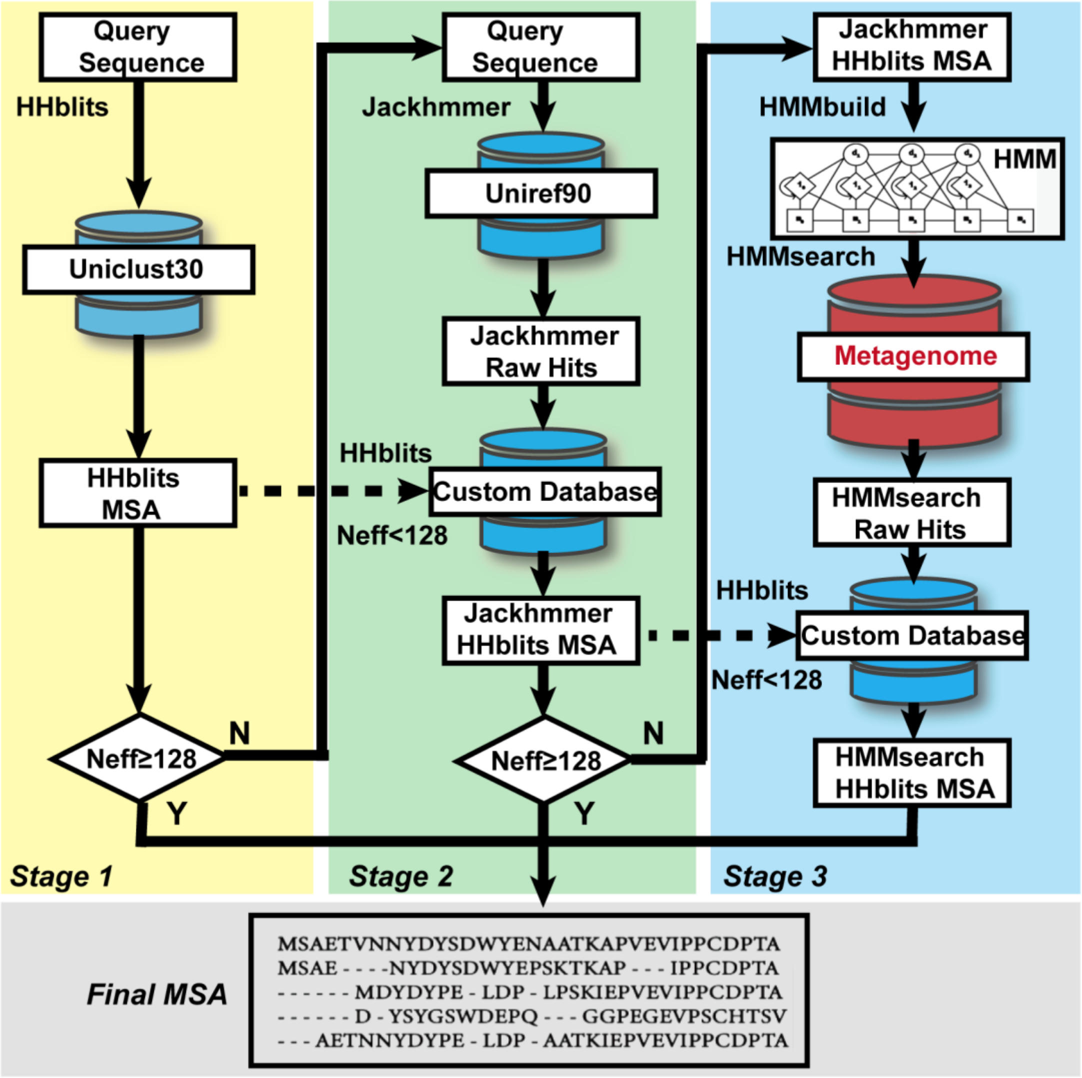

**Figure S6. DeepMSA pipeline for multiple sequence alignment generation.** The metagenome database in the third step can be the combination of four biomes (Fermentor, Gut, Lake and Soil) or each individual biome.

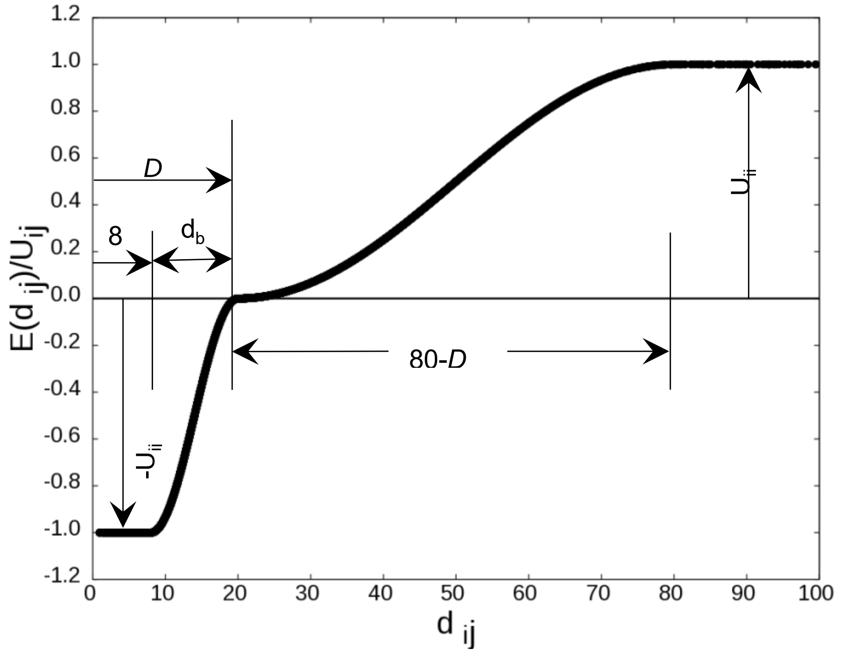

**Figure S7**: **A schematic of contact potential**, $E_{contact}(d_{ij})$, for a contacting residue pair $i$ and $j$ as defined in Eq. (S1). Here, $D$ is the protein length-dependent width of the first well and $U_{ij}$ is the depth of the energy potential that is proportional to the confidence score of the predicted contact between the residue pair $i$ and $j$. $d_{ij}$ is the $C_\beta$ distance between the residue pair.

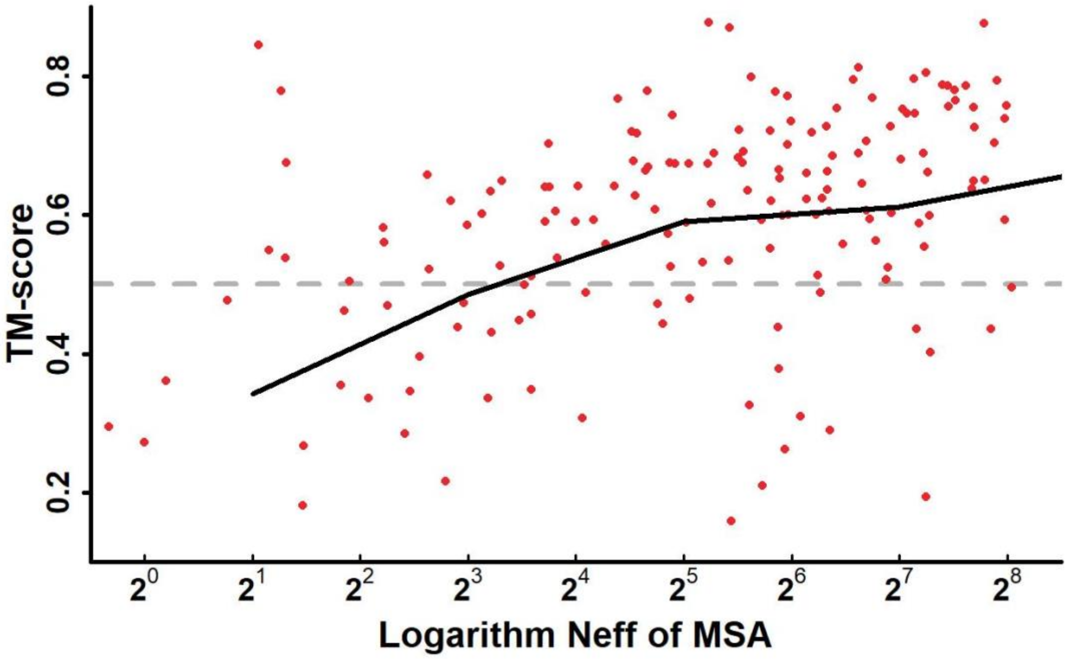

**Figure S8. TM-scores of the C-I-TASSER models from 168 proteins benchmark dataset for MSAs with different *Neff* values using a base of 2.** The black line represents the average TM-scores under each *Neff* bin with a bin width of two.

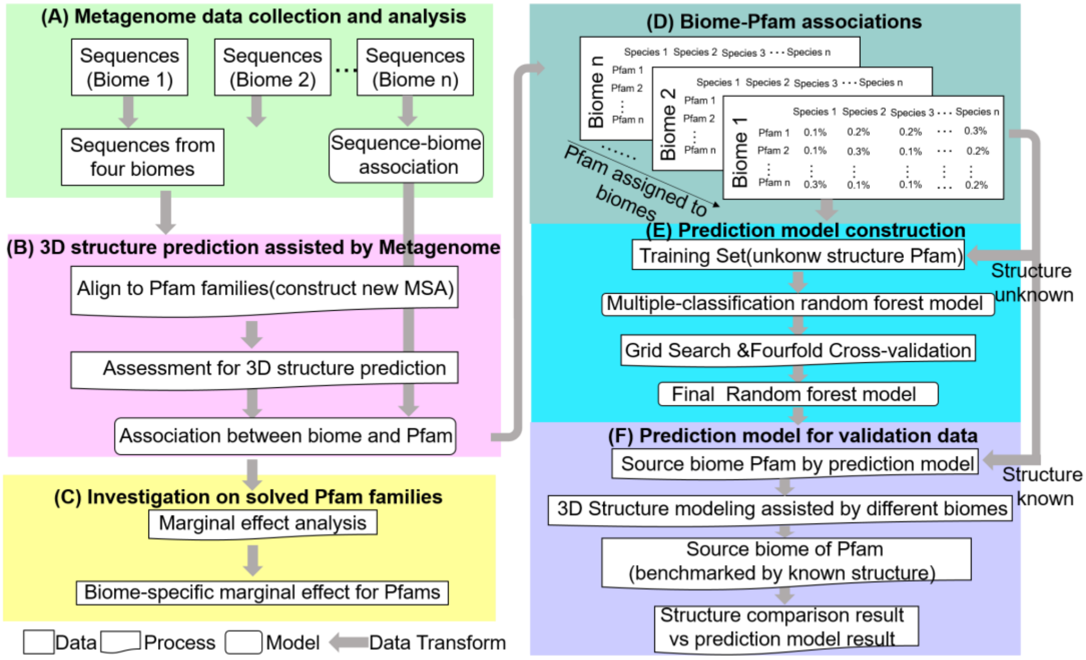

**Figure S9. Workflow for targeted MetaSource model construction.** (A) Sequences from different biomes were collected, and the biome-sequence associations are also organized. (B) New multiple sequence alignment (MSA) is constructed for Pfam families after search the homology sequences from different biomes. After the MSA is constructed, the *Neff*/C-socre were calculated to evaluate the quality of MSA. (C) The marginal effect is evaluated to quantify the effects of metagenome data from different biomes on Pfam families. (D) For each of the Pfam families, the normalized taxonomical composition was used as the feature. The biome with highest *Neff* score was used as the data label after supplementing the homology sequences from four biomes respectively. (E) The multiclass Random-Forest model construction. To find the best combination of model parameters, grid search was applied to exhaustive search over all parameter values and 20 cross-validation iterations. (F) The validation of MetaSource using Pfam families whose structure solved. Assisted by sequences from different biomes, the biome of which the structure that shared most similarity to the known structure is compared with the prediction result of MetaSource.

#### Supplementary Tables

**Table S1. Wilcox test results for differentiating each pair of two biomes based on species distribution.** Results shown are P-values of the Wilcox test.

| | **Fermentor** | **Gut** | **Lake** | **Soil** |
|---|---|---|---|---|
| **Fermentor** | | 2.53E-08 | 6.75E-10 | 8.23E-15 |
| **Gut** | 2.53E-08 | | 4.15E-15 | 7.23E-18 |
| **Lake** | 6.75E-10 | 4.15E-15 | | 6.28E-09 |
| **Soil** | 8.23E-15 | 7.23E-18 | 6.28E-09 | |

**Table S2. Summary of C-I-TASSER modeling results for 28 Pfam families which has solved experimental structure.** The comparison results for the solved protein to the C-I-TASSER model using TM-align and calculate the TM-score between the C-I-TASSER model and the map experimental structure.

| Target | *Neff* of MSA | PDB | TM-score | C-score |
|---|---|---|---|---|
| **PF04213** | 106.2 | 6JSB_A | 0.841 | 0.35 |
| **PF09139** | 34.5 | 6IG4_B | 0.763 | -1.16 |
| **PF03981** | 309.7 | 6RWT_A | 0.753 | -1.65 |
| **PF05914** | 19.1 | 6U42_4Q | 0.742 | -9.96 |
| **PF01803** | 28.6 | 6S9S_A | 0.717 | -0.99 |
| **PF04031** | 49.1 | 6OF2_A | 0.716 | -1.44 |
| **PF11704** | 22.8 | 6ULG_L | 0.684 | -0.57 |
| **PF12922** | 48.6 | 6QJ3_A | 0.672 | -1.45 |
| **PF18755** | 261.3 | 6PBD_B | 0.651 | -1.74 |
| **PF10785** | 33 | 6GCS_X | 0.639 | -0.6 |
| **PF03381** | 86.4 | 6PSY_E | 0.635 | -0.1 |
| **PF04317** | 30.5 | 6NZ4_A | 0.607 | -3.63 |
| **PF14687** | 24.1 | 6SGB_F6 | 0.6 | -1.31 |
| **PF15096** | 21 | 6R0X_E | 0.556 | -4.19 |
| **PF12017** | 87.8 | 6P5A_A | 0.465 | -3.43 |
| **PF13864** | 57.3 | 6U42_5S | 0.464 | -1.99 |
| **PF12357** | 70 | 6KZ8_B | 0.401 | -2.33 |
| **PF14260** | 101 | 6P1H_A | 0.381 | -5.47 |
| **PF04281** | 32.8 | 6JNF_C | 0.357 | -3.97 |
| **PF14636** | 16.1 | 6ULG_N | 0.322 | -2.51 |
| **PF07127** | 76.8 | 6U6G_A | 0.308 | -4.15 |
| **PF03963** | 196.5 | 6IEE_B | 0.272 | -3.66 |
| **PF12542** | 45.8 | 5YZG_X | 0.255 | -3.58 |
| **PF14960** | 18.4 | 6J5J_i | 0.249 | -3.07 |
| **PF14892** | 24.1 | 6U42_7H | 0.239 | -3.7 |
| **PF10172** | 20.6 | 6Q0R_E | 0.221 | -3.26 |
| **PF13868** | 44.2 | 6U42_4Y | 0.213 | -3.8 |
| **PF08648** | 70.8 | 6QX9_X | 0.182 | -3.96 |

**Table S3. The statistical result for GO annotations (level 3) which were only detected in single biome for the 964 Pfam families.** The numbers count for the GO entries that are only detected in a specific biome. The proportion of all entries detected in the corresponding biome under the specific top GO annotation was calculated.

| Biome | Biological Process | Molecular Function | Cellular Component |
|---|---|---|---|
| **Gut** | 21(30%) | 15(22%) | 18(25%) |
| **Lake** | 18(21%) | 11(25%) | 17(30%) |
| **Soil** | 42(33%) | 25(25%) | 44(30%) |
| **Fermentor** | 48(35%) | 18(25%) | 30(33%) |

**Table S4. Ten case studies for illustration of the Pfam-biome associations**. Ten Pfam families were selected based on the record in Pfam database and literature review. "Ferm" refers to "Fermentor"; "Data1" to "Uniref100"; "Data2" to "IMG+Uniref100"; "Data3" to "Tara Oceans+Uniref100"; "Data4" to "Metaclust+Uniref100"; "This work" to "Specific biom+Uniref100". Bold fonts highlight the best result for each target.

| **Pfam_ID** | **Source biome** | **Function** | **Accuracy** | **Neff for different databases** | | | | |
|---|---|---|---|---|---|---|---|---|
| | | | | **Data1** | **Data2** | **Data3** | **Data4** | **This work** |
| PF12652 | Ferm | CotJB protein; involed in the synthesis of spore coat related to anaerobic fermentation | Ferm:0.992 | 59.3 | 264 | 95.2 | 99.6058 | **Ferm:305.6** |
| PF06135 | Gut | IreB regulatory phosphoprotein, cephalosporin resistance | Gut:0.995 | 37.6 | **199.3** | 68.5 | 90.0926 | Gut:187.6 |
| PF07593 | Lake | ASPIC and UnbV | Lake:0.961 | 180.8 | 881.5 | 202.5 | 780.3 | **Lake:984.4** |
| PF09650 | Soil | Putative polyhydroxyalkanoic acid(PHA) system protein, detect in soil,Production of bioplastic | Soil:0.954 | 36.6 | 722.1 | 612.5 | 309.7 | **Soil:728.6** |
| PF13822 | | Acyl-CoA carboxylase epsilon subunit,involved in the biosynthesis of long-chain fatty acids | Soil:0.928 | 103.7 | 109.5 | 160.3 | 125.6 | **Soil:309.1** |
| PF04066 | | Multiple resistance and pH regulation protein F | Soil:0.936 | 168.1 | 844.8 | 240.502 | 525.5519 | **Soil:924.0** |
| PF09907 | | HigB_toxin, RelE-like toxic component of a toxin-antitoxin system | Soil:0.951 | 80 | 849.8 | 313.6 | 579.4 | **Soil:927.6** |
| PF09828 | | Chromate resistance exported protein | Soil:0.968 | 28.4 | 633.6 | 452.5 | 287.4 | **Soil:687.9** |
| PF05120 | | Gas vesicle protein G | Soil:0.986 | 32 | 336.8 | 69 | 178.9 | **Soil:487.5** |
| PF05425 | | Copper resistance protein D | Soil:0.961 | 257.2 | 389.5 | 364.2 | 726.1 | **Soil:807.8** |

**Table S5** are provided in an external Excel file "**Supplementary Tables.xlsx**".

**Table S6. The validation result of the MetaSource for the 204 Pfam families with solved structures.** MetaSource predicted the source biome which the queried Pfam could supplement the homology sequences. "ferm" refers to "fermentor".

| Pfam | Probability of source biome | | | | Predicted biome | Result based on | |
|---|---|---|---|---|---|---|---|
| | gut | lake | soil | ferm | | Neff | TM-score |
| **PF00284** | 2.13E-01 | 6.40E-02 | 6.44E-01 | 7.80E-02 | soil | soil | soil |
| **PF00631** | 8.27E-01 | 6.45E-02 | 1.32E-02 | 9.52E-02 | gut | gut | gut |
| **PF00647** | 1.26E-02 | 8.43E-02 | 8.22E-01 | 8.14E-02 | soil | soil | soil |
| **PF00658** | 1.09E-01 | 7.94E-02 | 7.46E-01 | 6.60E-02 | soil | soil | soil |
| **PF00737** | 6.40E-02 | 6.44E-01 | 2.13E-01 | 7.80E-02 | lake | lake | lake |
| **PF00827** | 1.13E-01 | 5.39E-02 | 7.75E-01 | 5.78E-02 | soil | ferm | ferm |
| **PF00833** | 1.13E-01 | 1.53E-01 | 6.91E-01 | 4.29E-02 | soil | lake | lake |
| **PF00838** | 1.26E-02 | 8.22E-01 | 8.43E-02 | 8.14E-02 | lake | lake | lake |
| **PF00853** | 3.88E-02 | 8.74E-01 | 4.22E-02 | 4.53E-02 | lake | lake | lake |
| **PF00960** | 1.38E-02 | 1.66E-01 | 6.57E-01 | 1.63E-01 | soil | soil | soil |
| **PF01049** | 8.74E-01 | 4.22E-02 | 3.88E-02 | 4.53E-02 | gut | gut | gut |
| **PF01111** | 1.09E-01 | 7.94E-02 | 7.46E-01 | 6.60E-02 | soil | lake | lake |
| **PF01115** | 1.26E-02 | 8.43E-02 | 8.22E-01 | 8.14E-02 | soil | ferm | ferm |
| **PF01125** | 1.09E-01 | 7.94E-02 | 7.46E-01 | 6.60E-02 | soil | soil | soil |
| **PF01140** | 7.81E-01 | 8.27E-02 | 2.06E-02 | 1.16E-01 | gut | gut | gut |
| **PF01191** | 1.13E-01 | 5.39E-02 | 7.75E-01 | 5.78E-02 | soil | soil | soil |
| **PF01194** | 1.09E-01 | 6.52E-01 | 1.73E-01 | 6.60E-02 | lake | lake | lake |
| **PF01200** | 1.09E-01 | 6.52E-01 | 1.73E-01 | 6.60E-02 | lake | lake | lake |
| **PF01213** | 3.22E-02 | 8.54E-01 | 6.81E-02 | 4.52E-02 | lake | lake | lake |
| **PF01214** | 7.46E-01 | 7.94E-02 | 1.09E-01 | 6.60E-02 | gut | gut | gut |
| **PF01247** | 1.09E-01 | 7.94E-02 | 7.46E-01 | 6.60E-02 | soil | soil | soil |
| **PF01267** | 1.03E-02 | 8.53E-01 | 5.58E-02 | 8.14E-02 | lake | soil | lake |
| **PF01278** | 1.04E-01 | 2.96E-02 | 8.04E-01 | 6.19E-02 | soil | soil | soil |
| **PF01320** | 1.64E-02 | 7.86E-01 | 4.23E-02 | 1.55E-01 | lake | lake | lake |
| **PF01340** | 4.03E-03 | 1.86E-02 | 9.51E-01 | 2.66E-02 | soil | soil | soil |
| **PF01356** | 1.16E-01 | 6.08E-01 | 1.09E-01 | 1.67E-01 | lake | lake | lake |
| **PF01603** | 7.20E-01 | 8.82E-02 | 1.12E-01 | 7.99E-02 | gut | gut | gut |
| **PF01716** | 2.13E-01 | 6.40E-02 | 6.44E-01 | 7.80E-02 | soil | soil | soil |
| **PF01780** | 1.09E-01 | 6.52E-01 | 1.73E-01 | 6.60E-02 | lake | lake | lake |
| **PF01793** | 9.01E-01 | 2.57E-02 | 2.69E-02 | 4.62E-02 | gut | gut | gut |
| **PF01815** | 1.61E-02 | 7.82E-02 | 8.17E-01 | 8.87E-02 | soil | soil | soil |
| **PF01821** | 5.13E-01 | 7.78E-02 | 3.58E-01 | 5.13E-02 | gut | gut | gut |
| **PF01828** | 8.85E-02 | 4.91E-02 | 1.77E-01 | 6.85E-01 | ferm | ferm | ferm |
| **PF01893** | 7.81E-01 | 8.27E-02 | 2.06E-02 | 1.16E-01 | gut | gut | gut |
| **PF01993** | 2.06E-02 | 8.27E-02 | 7.81E-01 | 1.16E-01 | soil | soil | soil |
| **PF02015** | 2.99E-02 | 7.56E-01 | 5.93E-02 | 1.55E-01 | lake | lake | lake |
| **PF02064** | 1.08E-02 | 9.20E-01 | 2.95E-02 | 3.97E-02 | lake | lake | lake |
| **PF02093** | 3.58E-01 | 7.78E-02 | 5.13E-01 | 5.13E-02 | soil | ferm | ferm |

| | | | | | | | |
|---|---|---|---|---|---|---|---|
| **PF02100** | 9.20E-01 | 2.95E-02 | 1.08E-02 | 3.97E-02 | gut | gut | gut |
| **PF02145** | 1.37E-02 | 8.13E-01 | 6.95E-02 | 1.04E-01 | lake | lake | lake |
| **PF02177** | 8.74E-01 | 4.22E-02 | 3.88E-02 | 4.53E-02 | gut | gut | gut |
| **PF02209** | 1.33E-02 | 8.52E-01 | 5.15E-02 | 8.34E-02 | lake | lake | lake |
| **PF02240** | 2.06E-02 | 8.27E-02 | 1.16E-01 | 7.81E-01 | ferm | ferm | ferm |
| **PF02253** | 0.00E+00 | 4.13E-03 | 9.57E-01 | 3.92E-02 | soil | ferm | ferm |
| **PF02271** | 1.07E-01 | 5.09E-02 | 7.77E-01 | 6.60E-02 | soil | ferm | ferm |
| **PF02284** | 9.20E-01 | 2.95E-02 | 1.08E-02 | 3.97E-02 | gut | gut | gut |
| **PF02289** | 2.06E-02 | 2.94E-01 | 4.90E-01 | 1.96E-01 | soil | soil | soil |
| **PF02312** | 3.88E-02 | 4.22E-02 | 8.74E-01 | 4.53E-02 | soil | soil | soil |
| **PF02315** | 1.93E-02 | 3.04E-01 | 6.04E-02 | 6.17E-01 | ferm | soil | soil |
| **PF02531** | 6.44E-01 | 6.40E-02 | 2.13E-01 | 7.80E-02 | gut | gut | gut |
| **PF02605** | 2.13E-01 | 6.44E-01 | 6.40E-02 | 7.80E-02 | lake | lake | lake |
| **PF02611** | 1.07E-01 | 2.62E-02 | 6.41E-01 | 2.26E-01 | soil | ferm | ferm |
| **PF02679** | 6.81E-02 | 8.30E-01 | 2.24E-02 | 7.93E-02 | lake | lake | lake |
| **PF02792** | 1.09E-01 | 7.46E-01 | 7.94E-02 | 6.60E-02 | lake | lake | lake |
| **PF02840** | 1.09E-01 | 7.94E-02 | 7.46E-01 | 6.60E-02 | soil | soil | soil |
| **PF02888** | 7.61E-01 | 7.96E-02 | 9.27E-02 | 6.68E-02 | gut | gut | gut |
| **PF02898** | 2.15E-01 | 4.74E-02 | 5.91E-01 | 1.46E-01 | soil | soil | soil |
| **PF02921** | 1.12E-01 | 8.82E-02 | 7.20E-01 | 7.99E-02 | soil | ferm | ferm |
| **PF02924** | 0.00E+00 | 4.13E-03 | 8.71E-01 | 1.25E-01 | soil | lake | ferm |
| **PF02963** | 1.42E-02 | 8.32E-01 | 3.96E-02 | 1.14E-01 | lake | lake | lake |
| **PF02974** | 7.94E-01 | 5.60E-02 | 9.33E-03 | 1.40E-01 | gut | gut | gut |
| **PF02975** | 8.27E-03 | 3.58E-02 | 8.96E-01 | 6.04E-02 | soil | ferm | ferm |
| **PF02979** | 2.12E-01 | 7.23E-01 | 4.02E-02 | 2.40E-02 | lake | lake | lake |
| **PF03013** | 6.51E-01 | 3.07E-01 | 4.49E-03 | 3.71E-02 | gut | gut | gut |
| **PF03095** | 7.46E-01 | 7.94E-02 | 1.09E-01 | 6.60E-02 | gut | gut | gut |
| **PF03110** | 2.13E-01 | 6.40E-02 | 6.44E-01 | 7.80E-02 | soil | soil | soil |
| **PF03126** | 1.09E-01 | 7.46E-01 | 7.94E-02 | 6.60E-02 | lake | lake | lake |
| **PF03288** | 3.63E-01 | 1.60E-02 | 4.77E-01 | 1.44E-01 | soil | soil | soil |
| **PF03411** | 1.09E-01 | 1.47E-01 | 6.25E-01 | 1.18E-01 | soil | ferm | ferm |
| **PF03416** | 1.09E-01 | 7.94E-02 | 7.46E-01 | 6.60E-02 | soil | ferm | ferm |
| **PF03502** | 9.51E-01 | 1.86E-02 | 4.03E-03 | 2.66E-02 | gut | gut | gut |
| **PF03660** | 1.97E-01 | 7.32E-02 | 6.64E-01 | 6.60E-02 | soil | ferm | ferm |
| **PF03735** | 1.10E-01 | 4.66E-02 | 7.76E-01 | 6.80E-02 | soil | soil | soil |
| **PF03829** | 2.54E-03 | 1.12E-01 | 7.61E-01 | 1.25E-01 | soil | soil | soil |
| **PF03870** | 1.09E-01 | 7.46E-01 | 7.94E-02 | 6.60E-02 | lake | lake | lake |
| **PF03887** | 7.13E-03 | 7.13E-01 | 3.47E-02 | 2.45E-01 | lake | lake | lake |
| **PF03925** | 1.08E-02 | 4.09E-02 | 8.74E-01 | 7.48E-02 | soil | soil | soil |
| **PF03974** | 7.74E-01 | 4.09E-02 | 1.08E-02 | 1.75E-01 | gut | gut | gut |
| **PF03997** | 1.09E-01 | 7.46E-01 | 7.94E-02 | 6.60E-02 | lake | lake | lake |
| **PF04008** | 1.03E-01 | 8.11E-01 | 1.38E-02 | 7.22E-02 | lake | lake | lake |
| **PF04038** | 2.06E-02 | 8.27E-02 | 7.81E-01 | 1.16E-01 | soil | soil | soil |
| **PF04062** | 1.07E-01 | 7.77E-01 | 5.09E-02 | 6.60E-02 | lake | lake | lake |
| **PF04098** | 0.00E+00 | 4.58E-02 | 9.04E-01 | 5.00E-02 | soil | soil | soil |
| **PF04216** | 1.00E-01 | 1.03E-01 | 4.87E-01 | 3.10E-01 | soil | soil | soil |

| | | | | | | | |
|---|---|---|---|---|---|---|---|
| **PF04269** | 1.08E-02 | 8.74E-01 | 4.09E-02 | 7.48E-02 | lake | lake | lake |
| **PF04270** | 7.53E-01 | 1.67E-01 | 1.61E-02 | 6.37E-02 | gut | gut | gut |
| **PF04300** | 7.59E-01 | 4.95E-02 | 1.03E-02 | 1.81E-01 | gut | gut | gut |
| **PF04362** | 3.97E-02 | 3.17E-01 | 5.93E-02 | 5.84E-01 | gut | ferm | ferm |
| **PF04386** | 1.04E-01 | 1.17E-01 | 1.18E-02 | 7.67E-01 | gut | ferm | ferm |
| **PF04433** | 7.20E-01 | 8.82E-02 | 1.12E-01 | 7.99E-02 | gut | gut | gut |
| **PF04502** | 1.09E-01 | 7.94E-02 | 6.60E-02 | 7.46E-01 | gut | ferm | ferm |
| **PF04591** | 8.17E-01 | 7.82E-02 | 1.61E-02 | 8.87E-02 | gut | gut | gut |
| **PF04621** | 3.55E-01 | 5.20E-01 | 7.58E-02 | 4.93E-02 | lake | lake | lake |
| **PF04721** | 3.88E-02 | 8.74E-01 | 4.22E-02 | 4.53E-02 | lake | lake | lake |
| **PF04729** | 1.09E-01 | 7.94E-02 | 6.60E-02 | 7.46E-01 | gut | ferm | ferm |
| **PF04739** | 1.09E-01 | 7.94E-02 | 6.60E-02 | 7.46E-01 | gut | ferm | ferm |
| **PF05005** | 1.35E-01 | 7.98E-01 | 3.73E-02 | 2.99E-02 | lake | lake | lake |
| **PF05023** | 8.52E-03 | 2.31E-02 | 8.15E-01 | 1.53E-01 | soil | soil | soil |
| **PF05026** | 7.51E-01 | 5.96E-02 | 1.09E-01 | 7.99E-02 | gut | gut | gut |
| **PF05153** | 1.87E-01 | 2.45E-02 | 1.60E-01 | 6.29E-01 | gut | ferm | ferm |
| **PF05247** | 8.27E-03 | 6.87E-01 | 2.44E-01 | 6.04E-02 | lake | lake | lake |
| **PF05280** | 1.08E-02 | 6.74E-01 | 2.41E-01 | 7.48E-02 | lake | lake | lake |
| **PF05303** | 1.03E-02 | 8.59E-01 | 4.95E-02 | 8.14E-02 | lake | lake | lake |
| **PF05321** | 1.61E-02 | 8.17E-01 | 7.82E-02 | 8.87E-02 | lake | lake | lake |
| **PF05354** | 2.13E-01 | 6.55E-01 | 6.07E-02 | 7.18E-02 | lake | lake | lake |
| **PF05370** | 7.81E-01 | 8.27E-02 | 2.06E-02 | 1.16E-01 | gut | gut | gut |
| **PF05551** | 2.16E-01 | 6.54E-01 | 9.57E-02 | 3.49E-02 | lake | lake | lake |
| **PF05854** | 7.81E-01 | 8.27E-02 | 2.06E-02 | 1.16E-01 | gut | gut | gut |
| **PF05856** | 1.09E-01 | 7.94E-02 | 7.46E-01 | 6.60E-02 | soil | soil | soil |
| **PF05870** | 8.47E-01 | 1.73E-02 | 3.10E-03 | 1.33E-01 | gut | gut | gut |
| **PF05983** | 1.12E-01 | 7.20E-01 | 8.82E-02 | 7.99E-02 | lake | lake | lake |
| **PF06141** | 2.29E-01 | 5.30E-01 | 1.55E-01 | 8.62E-02 | lake | lake | lake |
| **PF06154** | 1.61E-02 | 8.17E-01 | 7.82E-02 | 8.87E-02 | lake | lake | lake |
| **PF06175** | 1.11E-01 | 1.34E-01 | 6.78E-02 | 6.88E-01 | gut | ferm | ferm |
| **PF06304** | 2.10E-01 | 2.09E-01 | 5.67E-01 | 1.39E-02 | soil | soil | soil |
| **PF06384** | 6.67E-01 | 4.96E-02 | 1.07E-01 | 1.76E-01 | gut | gut | gut |
| **PF06400** | 3.58E-02 | 4.02E-02 | 8.81E-01 | 4.33E-02 | soil | soil | soil |
| **PF06438** | 1.08E-02 | 4.62E-02 | 9.06E-02 | 8.52E-01 | gut | ferm | ferm |
| **PF06456** | 3.88E-02 | 4.22E-02 | 4.53E-02 | 8.74E-01 | ferm | ferm | ferm |
| **PF06475** | 2.12E-01 | 7.06E-01 | 4.75E-02 | 3.42E-02 | lake | lake | lake |
| **PF06482** | 3.88E-02 | 4.22E-02 | 4.53E-02 | 8.74E-01 | ferm | ferm | ferm |
| **PF06557** | 2.06E-02 | 7.81E-01 | 8.27E-02 | 1.16E-01 | lake | lake | lake |
| **PF06684** | 3.67E-01 | 2.17E-01 | 2.04E-01 | 2.12E-01 | gut | gut | gut |
| **PF06844** | 1.04E-01 | 1.19E-01 | 7.41E-01 | 3.64E-02 | soil | ferm | ferm |
| **PF06870** | 7.51E-01 | 5.96E-02 | 1.09E-01 | 7.99E-02 | gut | gut | gut |
| **PF07072** | 1.08E-02 | 3.40E-01 | 5.90E-01 | 6.00E-02 | soil | ferm | ferm |
| **PF07152** | 1.04E-01 | 6.67E-01 | 1.17E-01 | 1.12E-01 | lake | lake | lake |
| **PF07262** | 1.99E-02 | 8.22E-01 | 8.37E-02 | 7.43E-02 | lake | lake | lake |
| **PF07352** | 2.38E-03 | 5.29E-01 | 4.12E-01 | 5.71E-02 | lake | ferm | ferm |
| **PF07361** | 1.08E-02 | 3.97E-02 | 8.90E-01 | 6.00E-02 | soil | soil | soil |

| | | | | | | | |
|---|---|---|---|---|---|---|---|
| **PF07408** | 6.69E-01 | 5.97E-02 | 1.52E-01 | 1.19E-01 | gut | gut | gut |
| **PF07460** | 1.19E-01 | 5.82E-02 | 5.74E-01 | 2.48E-01 | soil | soil | soil |
| **PF07472** | 1.82E-02 | 5.41E-02 | 1.16E-01 | 8.12E-01 | gut | ferm | ferm |
| **PF07682** | 2.32E-01 | 5.97E-02 | 5.89E-01 | 1.19E-01 | soil | soil | soil |
| **PF07828** | 1.61E-02 | 7.82E-02 | 8.17E-01 | 8.87E-02 | soil | soil | soil |
| **PF08000** | 2.04E-01 | 4.49E-03 | 2.41E-01 | 5.50E-01 | gut | ferm | ferm |
| **PF08127** | 8.91E-02 | 7.82E-01 | 7.26E-02 | 5.64E-02 | lake | lake | lake |
| **PF08208** | 2.98E-02 | 3.95E-02 | 4.52E-02 | 8.85E-01 | gut | ferm | ferm |
| **PF08536** | 2.13E-01 | 6.40E-02 | 6.44E-01 | 7.80E-02 | soil | soil | soil |
| **PF08714** | 9.51E-03 | 2.60E-01 | 4.46E-01 | 2.84E-01 | soil | soil | soil |
| **PF08773** | 2.20E-01 | 3.84E-02 | 2.97E-02 | 7.12E-01 | gut | ferm | ferm |
| **PF08804** | 1.08E-02 | 8.74E-01 | 4.09E-02 | 7.48E-02 | lake | lake | lake |
| **PF08814** | 2.06E-02 | 8.27E-02 | 1.16E-01 | 7.81E-01 | gut | ferm | ferm |
| **PF08854** | 6.31E-01 | 6.90E-02 | 2.14E-01 | 8.64E-02 | gut | gut | gut |
| **PF08869** | 6.29E-01 | 1.07E-01 | 2.80E-02 | 2.36E-01 | gut | gut | gut |
| **PF08883** | 3.20E-02 | 3.52E-02 | 8.51E-01 | 8.14E-02 | soil | soil | soil |
| **PF08931** | 2.06E-02 | 7.81E-01 | 8.27E-02 | 1.16E-01 | lake | lake | lake |
| **PF08941** | 3.58E-02 | 4.02E-02 | 8.81E-01 | 4.33E-02 | soil | soil | soil |
| **PF08958** | 5.89E-01 | 5.97E-02 | 2.32E-01 | 1.19E-01 | gut | gut | gut |
| **PF08963** | 5.89E-01 | 5.97E-02 | 2.32E-01 | 1.19E-01 | gut | gut | gut |
| **PF08968** | 5.89E-01 | 5.97E-02 | 2.32E-01 | 1.19E-01 | gut | gut | gut |
| **PF08974** | 1.09E-01 | 5.47E-02 | 1.26E-01 | 7.10E-01 | gut | ferm | ferm |
| **PF08992** | 8.57E-02 | 7.22E-02 | 1.47E-01 | 6.95E-01 | ferm | ferm | ferm |
| **PF09001** | 7.81E-01 | 8.27E-02 | 2.06E-02 | 1.16E-01 | gut | gut | gut |
| **PF09009** | 3.20E-02 | 3.81E-02 | 1.95E-01 | 7.35E-01 | ferm | ferm | ferm |
| **PF09015** | 9.79E-03 | 4.47E-02 | 5.09E-02 | 8.95E-01 | ferm | ferm | ferm |
| **PF09021** | 2.13E-01 | 1.52E-01 | 1.60E-01 | 4.74E-01 | ferm | ferm | ferm |
| **PF09028** | 1.82E-02 | 7.12E-01 | 1.54E-01 | 1.16E-01 | lake | lake | lake |
| **PF09044** | 1.37E-02 | 4.67E-02 | 8.61E-01 | 7.85E-02 | soil | soil | soil |
| **PF09056** | 1.49E-02 | 6.23E-02 | 7.96E-01 | 1.27E-01 | soil | soil | soil |
| **PF09059** | 1.08E-02 | 4.09E-02 | 8.74E-01 | 7.48E-02 | soil | soil | soil |
| **PF09078** | 1.08E-02 | 2.41E-01 | 7.48E-02 | 6.74E-01 | ferm | ferm | ferm |
| **PF09082** | 2.06E-02 | 8.15E-02 | 2.01E-01 | 6.97E-01 | ferm | ferm | ferm |
| **PF09143** | 1.37E-02 | 4.96E-02 | 8.45E-01 | 9.18E-02 | soil | soil | soil |
| **PF09160** | 1.61E-02 | 7.82E-02 | 8.17E-01 | 8.87E-02 | soil | soil | soil |
| **PF09194** | 1.82E-02 | 5.41E-02 | 8.09E-01 | 1.19E-01 | soil | soil | soil |
| **PF09203** | 8.69E-01 | 6.43E-02 | 1.65E-02 | 5.06E-02 | gut | gut | gut |
| **PF09204** | 8.12E-01 | 3.46E-02 | 8.27E-03 | 1.46E-01 | gut | gut | gut |
| **PF09208** | 1.47E-01 | 7.25E-01 | 2.24E-02 | 1.06E-01 | lake | lake | lake |
| **PF09218** | 7.81E-01 | 8.27E-02 | 2.06E-02 | 1.16E-01 | gut | gut | gut |
| **PF09221** | 9.66E-02 | 5.97E-02 | 7.47E-01 | 9.71E-02 | soil | lake | soil |
| **PF09223** | 1.35E-01 | 1.89E-01 | 6.04E-01 | 7.18E-02 | soil | soil | soil |
| **PF09225** | 2.10E-01 | 1.06E-02 | 4.55E-01 | 3.25E-01 | soil | soil | soil |
| **PF09226** | 1.82E-02 | 5.41E-02 | 8.09E-01 | 1.19E-01 | soil | soil | soil |
| **PF09233** | 8.51E-01 | 5.43E-02 | 9.25E-03 | 8.51E-02 | gut | gut | gut |
| **PF09391** | 8.25E-01 | 4.80E-02 | 1.03E-01 | 2.46E-02 | gut | gut | gut |

| | | | | | | | |
|---|---|---|---|---|---|---|---|
| **PF09392** | 9.51E-01 | 1.86E-02 | 4.03E-03 | 2.66E-02 | gut | gut | gut |
| **PF09393** | 2.13E-01 | 1.60E-01 | 5.14E-01 | 1.13E-01 | soil | soil | soil |
| **PF09412** | 3.83E-02 | 3.72E-02 | 3.69E-02 | 8.88E-01 | ferm | ferm | ferm |
| **PF09449** | 2.06E-02 | 8.27E-02 | 7.81E-01 | 1.16E-01 | soil | gut | gut |
| **PF09628** | 2.32E-01 | 5.89E-01 | 5.97E-02 | 1.19E-01 | lake | lake | lake |
| **PF09642** | 2.32E-01 | 5.89E-01 | 5.97E-02 | 1.19E-01 | lake | lake | lake |
| **PF10054** | 1.09E-01 | 8.22E-01 | 4.93E-02 | 2.03E-02 | lake | lake | lake |
| **PF10120** | 1.11E-01 | 7.34E-01 | 6.89E-02 | 8.57E-02 | lake | lake | lake |
| **PF10634** | 2.54E-03 | 3.82E-01 | 6.80E-02 | 5.48E-01 | ferm | ferm | ferm |
| **PF11102** | 9.48E-01 | 1.86E-02 | 4.03E-03 | 2.97E-02 | gut | ferm | ferm |
| **PF11419** | 2.06E-02 | 8.27E-02 | 7.81E-01 | 1.16E-01 | soil | soil | soil |
| **PF11428** | 2.32E-01 | 5.97E-02 | 5.89E-01 | 1.19E-01 | soil | soil | soil |
| **PF11429** | 2.15E-01 | 3.40E-02 | 5.84E-01 | 1.67E-01 | soil | gut | gut |
| **PF11432** | 2.06E-02 | 7.81E-01 | 8.27E-02 | 1.16E-01 | lake | lake | lake |
| **PF11436** | 2.32E-01 | 5.97E-02 | 5.89E-01 | 1.19E-01 | soil | gut | gut |
| **PF11497** | 2.06E-02 | 8.27E-02 | 7.81E-01 | 1.16E-01 | soil | soil | soil |
| **PF11644** | 2.06E-02 | 7.81E-01 | 8.27E-02 | 1.16E-01 | lake | lake | lake |
| **PF11708** | 1.12E-01 | 8.82E-02 | 7.20E-01 | 7.99E-02 | soil | gut | gut |
| **PF11724** | 3.27E-01 | 1.51E-02 | 5.63E-01 | 9.54E-02 | soil | soil | soil |
| **PF12106** | 2.38E-01 | 5.63E-01 | 9.38E-02 | 1.05E-01 | lake | lake | lake |
| **PF12134** | 1.09E-01 | 7.94E-02 | 7.46E-01 | 6.60E-02 | soil | soil | soil |
| **PF12924** | 3.88E-02 | 4.22E-02 | 8.74E-01 | 4.53E-02 | soil | soil | soil |
| **PF14511** | 1.68E-02 | 1.32E-01 | 7.70E-01 | 8.07E-02 | soil | gut | gut |
| **PF14562** | 2.06E-02 | 8.27E-02 | 7.81E-01 | 1.16E-01 | soil | ferm | ferm |
| **PF15009** | 2.20E-01 | 3.84E-02 | 7.12E-01 | 2.97E-02 | soil | gut | gut |
| **PF18484** | 1.08E-02 | 3.41E-01 | 5.74E-01 | 7.48E-02 | soil | gut | gut |
| **PF18681** | 2.32E-01 | 5.97E-02 | 5.89E-01 | 1.19E-01 | soil | soil | soil |
| **PF18882** | 1.11E-01 | 6.89E-02 | 7.34E-01 | 8.57E-02 | soil | soil | soil |

**References**

1. Mitchell, A.L., Scheremetjew, M., Denise, H., Potter, S., Tarkowska, A., Qureshi, M., Salazar, G.A., Pesseat, S., Boland, M.A., Hunter, F.M.I. *et al.* (2018) EBI Metagenomics in 2017: enriching the analysis of microbial communities, from sequence reads to assemblies. *Nucleic Acids Res*, **46**, D726-D735.
2. Eddy, S.R. (1998) Profile hidden Markov models. *Bioinformatics*, **14**, 755-763.
3. Zheng, W., Zhang, C., Wuyun, Q., Pearce, R., Li, Y. and Zhang, Y. (2019) LOMETS2: improved meta-threading server for fold-recognition and structure-based function annotation for distant-homology proteins. *Nucleic Acids Res*, **47**, W429-W436.
4. Zhang, C., Zheng, W., Mortuza, S.M., Li, Y. and Zhang, Y. (2020) DeepMSA: constructing deep multiple sequence alignment to improve contact prediction and fold-recognition for distant-homology proteins. *Bioinformatics*, **36**, 2105-2112.
5. Remmert, M., Biegert, A., Hauser, A. and Söding, J. (2012) HHblits: lightning-fast iterative protein sequence searching by HMM-HMM alignment. *Nature Methods*, **9**, 173-175.
6. Mirdita, M., von den Driesch, L., Galiez, C., Martin, M.J., Söding, J. and Steinegger, M. (2017) Uniclust databases of clustered and deeply annotated protein sequences and alignments. *Nucleic Acids Research*, **45**, D170-D176.
7. Suzek, B.E., Wang, Y., Huang, H., McGarvey, P.B., Wu, C.H. and the UniProt, C. (2014) UniRef clusters: a comprehensive and scalable alternative for improving sequence similarity searches. *Bioinformatics*, **31**, 926-932.
8. Li, Y., Zhang, C., Bell, E.W., Zheng, W., Zhou, X., Yu, D.J. and Zhang, Y. (2021) Deducing high-accuracy protein contact-maps from a triplet of coevolutionary matrices through deep residual convolutional networks. *PLoS Comput Biol*, doi: https://doi.org/10.1371/journal.pcbi.1008865.
9. Li, Y., Zhang, C., Bell, E.W., Yu, D.J. and Zhang, Y. (2019) Ensembling multiple raw coevolutionary features with deep residual neural networks for contact-map prediction in CASP13. *Proteins*, **87**, 1082-1091.
10. He, B., Mortuza, S.M., Wang, Y., Shen, H.B. and Zhang, Y. (2017) NeBcon: protein contact map prediction using neural network training coupled with naive Bayes classifiers. *Bioinformatics*, **33**, 2296-2306.
11. Li, Y., Hu, J., Zhang, C., Yu, D.-J. and Zhang, Y. (2019) ResPRE: high-accuracy protein contact prediction by coupling precision matrix with deep residual neural networks. *Bioinformatics*, btz291.
12. Zheng, W., Li, Y., Zhang, C., Pearce, R., Mortuza, S.M. and Zhang, Y. (2019) Deep-learning contact-map guided protein structure prediction in CASP13. *Proteins*, **87**, 1149-1164.
13. Zheng, W., Wuyun, Q., Li, Y., Mortuza, S.M., Zhang, C., Pearce, R., Ruan, J. and Zhang, Y. (2019) Detecting distant-homology protein structures by aligning deep neural-network based contact maps. *PLoS Comput Biol*, **15**, e1007411.
14. Zhao, Y., Tagami, A., Dobele, G., Lindstrom, M.E. and Sevastyanova, O. (2019) The Impact of Lignin Structural Diversity on Performance of Cellulose Nanofiber (CNF)-Starch Composite Films. *Polymers (Basel)*, **11**.
15. Xu, D., Jaroszewski, L., Li, Z. and Godzik, A. (2014) FFAS-3D: improving fold recognition by including optimized structural features and template re-ranking. *Bioinformatics*, **30**, 660-667.

16. Soding, J. (2005) Protein homology detection by HMM-HMM comparison. *Bioinformatics*, **21**, 951-960.
17. Wu, S. and Zhang, Y. (2008) MUSTER: Improving protein sequence profile-profile alignments by using multiple sources of structure information. *Proteins*, **72**, 547-556.
18. Yang, J., Yan, R., Roy, A., Xu, D., Poisson, J. and Zhang, Y. (2015) The I-TASSER Suite: protein structure and function prediction. *Nat Methods*, **12**, 7-8.
19. Wu, S. and Zhang, Y. (2007) LOMETS: A local meta-threading-server for protein structure prediction. *Nucl. Acids. Res.*, **35**, 3375-3382.
20. Kim, D., Xu, D., Guo, J.T., Ellrott, K. and Xu, Y. (2003) PROSPECT II: protein structure prediction program for genome-scale applications. *Protein Eng*, **16**, 641-650.
21. Hughes, C.S., Foehr, S., Garfield, D.A., Furlong, E.E., Steinmetz, L.M. and Krijgsveld, J. (2014) Ultrasensitive proteome analysis using paramagnetic bead technology. *Mol Syst Biol*, **10**, 757.
22. Yang, Y., Faraggi, E., Zhao, H. and Zhou, Y. (2011) Improving protein fold recognition and template-based modeling by employing probabilistic-based matching between predicted one-dimensional structural properties of query and corresponding native properties of templates. *Bioinformatics*, **27**, 2076-2082.
23. Zhang, Y. and Skolnick, J. (2004) SPICKER: A clustering approach to identify near-native protein folds. *Journal of Computational Chemistry*, **25**, 865-871.
24. Zhang, J., Liang, Y. and Zhang, Y. (2011) Atomic-level protein structure refinement using fragment-guided molecular dynamics conformation sampling. *Structure*, **19**, 1784-1795.
25. Ovchinnikov, S., Park, H., Varghese, N., Huang, P.S., Pavlopoulos, G.A., Kim, D.E., Kamisetty, H., Kyrpides, N.C. and Baker, D. (2017) Protein structure determination using metagenome sequence data. *Science*, **355**, 294-298.
26. Wang, Y., Shi, Q., Yang, P., Zhang, C., Mortuza, S.M., Xue, Z., Ning, K. and Zhang, Y. (2019) Fueling ab initio folding with marine metagenomics enables structure and function predictions of new protein families. *Genome Biol*, **20**, 229.
27. Pfeifer, F. (2012) Distribution, formation and regulation of gas vesicles. *Nat Rev Microbiol*, **10**, 705-715.
28. van Keulen, G., Hopwood, D.A., Dijkhuizen, L. and Sawers, R.G. (2005) Gas vesicles in actinomycetes: old buoys in novel habitats? *Trends Microbiol*, **13**, 350-354.
29. Cheng, C., Bao, T. and Yang, S.T. (2019) Engineering Clostridium for improved solvent production: recent progress and perspective. *Appl Microbiol Biotechnol*, **103**, 5549-5566.
30. Bourassa, D.V., Kannenberg, E.L., Sherrier, D.J., Buhr, R.J. and Carlson, R.W. (2017) The Lipopolysaccharide Lipid A Long-Chain Fatty Acid Is Important for Rhizobium leguminosarum Growth and Stress Adaptation in Free-Living and Nodule Environments. *Mol Plant Microbe Interact*, **30**, 161-175.
31. Cornu, J.Y., Huguenot, D., Jezequel, K., Lollier, M. and Lebeau, T. (2017) Bioremediation of copper-contaminated soils by bacteria. *World J Microbiol Biotechnol*, **33**, 26.
32. Tamindzija, D., Chromikova, Z., Spaic, A., Barak, I., Bernier-Latmani, R. and Radnovic, D. (2019) Chromate tolerance and removal of bacterial strains isolated from uncontaminated and chromium-polluted environments. *World J Microbiol Biotechnol*, **35**, 56.
33. Cheng, J. and Charles, T.C. (2016) Novel polyhydroxyalkanoate copolymers produced in Pseudomonas putida by metagenomic polyhydroxyalkanoate synthases. *Appl Microbiol*

*Biotechnol*, **100**, 7611-7627.
